# Compensating Cortical Thickness for Cortical Folding-Related Variation

**DOI:** 10.1101/2025.05.03.651968

**Authors:** Nagehan Demirci, Timothy S. Coalson, Maria A. Holland, David C. Van Essen, Matthew F. Glasser

## Abstract

Cortical thickness is a widely used biomarker of brain morphology and health, yet it is dependent on local cortical folding. Because gyral crowns are consistently thicker than sulcal fundi and cortical folds vary widely across individuals, these fluctuations introduce unmodeled nuisance variance that can obscure meaningful biological effects of interest. Previous global methods of folding compensation incompletely compensate for folding effects on cortical thickness. Spatial smoothing is commonly used to reduce these effects in the literature, but this markedly degrades spatial localization precision. To address these limitations, we developed a novel method for folding-compensated cortical thickness estimation that uses nonlinear local multiple regression with five folding measures to model and more completely remove folding-related variance from cortical thickness. This approach estimates what cortical thickness would have been in the absence of folding, yielding a more biologically interpretable measure of cortical architecture. We applied this new approach to data from the Young Adult Human Connectome Project (HCP-YA) and Aging Human Connectome Project (HCA), demonstrating substantial reductions in intra-areal and inter-individual variability, substantially increasing standardized effect sizes of age on cortical thickness (41% increase) while preserving neurobiologically expected patterns, and avoiding the loss of spatial precision that occurs with the spatial smoothing that has traditionally been used in the literature. The method has been integrated into the HCP pipelines, facilitating its widespread use. By attenuating folding-induced variability, this technique enhances cortical thickness as a structural phenotype and may support more accurate cortical parcellation, longitudinal tracking, and biomarker discovery in brain health and disease.

## 1. Introduction

The human cerebral cortex is the highly convoluted outer layer of the forebrain that is responsible for a wide range of complex cognitive, sensory, and motor functions and is organized into distinct cortical areas and functional networks. To accommodate a large surface area within a limited intracranial volume, in many species the cortex undergoes extensive folding during development (Akula et al., 2023; Fernández et al., 2016; Van Essen, 2023; Welker, 1990). In most large-brained mammals, such as humans, the cortex is highly folded, whereas in smaller-brained species, such as rodents and prosimians, it remains largely smooth with minimal or no folding (Demirci et al., 2023a; Heuer et al., 2019; Van Essen et al., 2019). This folding gives rise to a characteristic pattern of gyri (ridges) and sulci (grooves), which increases cortical surface area while facilitating compact neural wiring (Van Essen, 1997) and optimizing the anatomical and functional organization of the cortex.

The thickness of the cerebral cortex, a critical indicator of cortical architecture, is consistently influenced by cortical folding patterns. This finding was first systematically described by Dutch neuroanatomist Siegfried Thomas Bok (1892-1964) in 1929 (Bok, 1929; Consolini et al., 2022). He noted that cortex is thicker in a convex curve (gyrus) and thinner in a concave curve (sulcal fundus) and that folding tends to preserve the volume of each cortical layer, as described in his seminal *equi-volume principle.* Cortical areas also differ in their average thicknesses due to differences in their microstructural architecture (i.e., differences in the thickness of their constituent cortical layers). In lissencephalic species such as the marmoset, most of the variability in cortical thickness across the cerebral cortex is attributable to inter-areal architectural differences. In more gyrencephalic species, such as macaques, apes, and human (Hayashi et al., 2021), thickness variations reflect a mixture of folding-related and intrinsic architectonic differences. Architectural variability in cortical thickness of gyrencephalic primates is particularly evident within the central sulcus, where the anterior bank (motor cortex) is much thicker than the posterior bank (somatosensory cortex) despite both banks being relatively flat (Biega et al., 2006; Meyer et al., 1996). Bok recognized this distinction between architectural and folding effects on cortical thickness and proposed that the effects due to folding are not functionally meaningful, in opposition to the prevailing views of his contemporaries. He emphasized the need to correct cortical thickness measures to account for these folding effects. Bok anticipated that *"[this] critical correction will be an enormous undertaking,"* recognizing the methodological challenges of his time.

Recent advances in MRI-based brain imaging and surface-based analysis have made it possible to address Bok’s century-old challenge, enabling compensation for folding-induced effects on cortical thickness estimates. A key advance was the development of FreeSurfer software for accurately reconstructing the pial and white surfaces and automatically computing cortical thickness (Fischl and Dale, 2000). In an early effort to computationally compensate for folding effects in cortical thickness and thereby estimate what the cortical thickness would have been in the absence of cortical folding, Sigalovsky et al., (2006) showed a substantial correlation (𝑟 = −0.56) between cortical thickness and mean cortical curvature in human temporal cortex. They then linearly regressed mean cortical curvature out of cortical thickness within the temporal lobes and attempted to threshold the residual cortical thickness to define cortical areas. Unfortunately, their analysis failed to reveal distinct cortical areas. Subsequently, a global curvature regression method was applied to the entire cortex and, together with myelin maps based on the ratio of T1- weighted / T2-weighted MRI images, contributed to the identification of dozens of cortical areas (Glasser and Van Essen, 2011). Importantly, cortical areal boundaries were delineated using spatial gradients computed on cortical surface meshes for both measures, rather than the thresholding approach (Sigalovsky et al., 2006). These gradients represent an objective, observer-independent measure of putative areal boundaries, inspired by analogous histological approaches (Schleicher et al., 1999) and are far better than thresholding for defining areal borders. The gradient-based approach was then extended to task-based and resting state fMRI and, together with T1w/T2w myelin maps and globally compensated cortical thickness maps, was used to parcellate each human cerebral hemisphere into 180 cortical areas (Glasser et al., 2016).

Global regression of mean curvature from cortical thickness maps leaves substantial folding-related variation unaccounted for, however. This is because the influence of folding on cortical thickness depends on what the underlying, *intrinsic* thickness would have been in the absence of folding—precisely what we aim to estimate—which varies regionally across the cortical sheet. As a result, applying a single global regression coefficient often underfits the regionally dependent effects of folding on cortical thickness. Additionally, mean curvature is an incomplete description of the geometric properties of cortical folding, which can also be described by various other folding measures, i.e., principal curvatures, Gaussian curvature, shape index, and curvedness. Finally, a linear regression only addresses first-order linear effects, whereas the effects of folding on cortical thickness are nonlinear (Demirci and Holland, 2024). Here we address these limitations by introducing a compensated cortical thickness metric based on local nonlinear multiple regression, which aims to reduce the influence of folding on cortical thickness. Thus, our approach aims to preserve neurobiologically meaningful thickness variations, such as those between motor and sensory cortex (Triarhou, 2007) or the atrophy that occurs with brain aging (Lemaitre et al., 2012; Lindroth et al., 2019; Salat et al., 2004), while substantially attenuating variations in observed cortical thickness due to cortical folding that vary widely across individuals and are not related to cortical areas or other effects of interest. This approach is similar in spirit to denoising approaches that we have applied in functional MRI, where the aim is to separate genuine neural fluctuations in the BOLD fMRI signal from nuisance fluctuations due to head motion or subject physiology (Glasser et al., 2018). To the extent that the influence of cortical folding is reduced while the sharpness of genuine transitions in cortical thickness or other effects such as atrophy are preserved, this approach can enable more robust comparisons of cortical thickness across individuals and cortical areas.

In previous human neuroimaging studies of cortical thickness (Demirci et al., 2023b; Duan et al., 2020; Gilmore et al., 2020; Gordon et al., 2015; Mueller et al., 2013) nuisance inter-individual variability in human cortical folding patterns has been addressed primarily through two approaches: (i) averaging thickness measurements within parcels defined by gyral-sulcal folds (Kim et al., 2012; Lyall et al., 2015; Narayana et al., 2013; Rosas et al., 2002; Tahedl, 2020) or by well-defined cortical areas (Alvarez et al., 2019; Glasser et al., 2016) or (ii) applying substantial spatial smoothing to cortical thickness maps (Chung et al., 2005; Doyle-Thomas et al., 2013; Hurtz et al., 2014; Hutton et al., 2008; Kim et al., 2012; Kuperberg et al., 2003; Lerch et al., 2005; Misaki et al., 2012; Nam et al., 2015; Rosas et al., 2002; Salat et al., 2004). However, areal averaging does not fully eliminate folding nuisance variability across individuals, thereby reducing statistical sensitivity. Further, heavy smoothing (mean 24 mm full-width-half-maximum (FWHM), see Supplementary Table 1) markedly diminishes the spatial specificity of findings, even when performed on surface meshes (Coalson et al., 2018).

This study introduces our new approach to compensating cortical thickness for folding effects and applies it to populations from the Human Connectome Project (HCP) (Bookheimer et al., 2019; Van Essen et al., 2013), focusing on young adults (22–35 years) and aging populations (36–100+ years). Unlike traditional methods that rely on high levels of smoothing to minimize nuisance folding variations, our compensation method preserves spatial localization precision while more completely accounting for folding-induced variation across individuals than prior global compensation methods. Importantly our approach does not alter the meaningful signal of interest and can still be averaged within well-defined cortical areas. Our work builds on a strong historical foundation and addresses longstanding questions about cortical folding and its impact on thickness. To enable other researchers to use this useful measure of cortical thickness, we have integrated this method into the publicly available HCP pipelines (Glasser et al., 2013), and our method will be generated both by default in new HCP Pipeline Structural Preprocessing runs and can also be applied to existing HCP Pipeline outputs without requiring rerunning of the full HCP Structural Preprocessing pipelines.

## 2. Methods

### 2.1 Datasets

We computed the folding-compensated cortical thickness for two independent datasets, the HCP Young Adult (HCP-YA) and HCP Aging (HCA) cohorts. The HCP-YA dataset includes 1071 healthy young adults (twins and non-twin siblings) aged between 22-35 years (485M, 586F) and the HCP Aging dataset includes 1798 individuals’ sessions aged between 36-103 years (520M, 681F; 597 repeat visits for longitudinal scanning). All human participants provided informed consent, and the HCP studies were approved by the Washington University Institutional Review Board (Bookheimer et al., 2019; Van Essen et al., 2013). Both datasets include high-quality T1w and T2w structural images, which are vital for generating accurate white and pial surface meshes of the cerebral cortex. For the HCP-YA dataset, the structural images were acquired at 0.7 mm isotropic resolution using a custom 3T Siemens Skyra and a 32-channel head coil, enabling accurate individual maps of myelin and cortical thickness (for details of structural image acquisition, see Glasser et al., (2022, 2013); Uğurbil et al., (2013). The HCA dataset acquisition protocol is similar to that used for the HCP-YA dataset, but the spatial resolution was adjusted to 0.8 mm isotropic, and scans were acquired on a Siemens 3T Prisma. For further details on the HCA dataset acquisition, see Glasser et al., (2022); Harms et al., (2018). Both datasets follow the HCP-style approach (Elam et al., 2021; Glasser et al., 2016) for structural MRI, acquiring images with voxels half the minimum cortical thickness (∼1.6 mm) or less in contrast to the traditional 1 mm T1w image resolution so as to improve the accuracy of cortical surface placement and thereby measures of cortical thickness and myelin. These data were processed through the HCP Pipelines (Glasser et al., 2013), which included the generation of white and pial surface meshes, cortical thickness measurement with FreeSurfer (Fischl, 2012), and the incorporation of T2-weighted imaging to refine pial surface placement. Additionally, cortical surface meshes were aligned across individuals using MSMAll areal-feature-based registration (Robinson et al., 2018) for optimal correspondence to the HCP’s multimodal cortical parcellation version 1.0 (Glasser et al., 2016). T1w/T2w myelin maps (with transmit field correction) and gradients and resting state functional connectivity gradients for comparison with these compensated cortical thickness gradient maps were generated as previously described (Glasser et al., 2022, 2019, 2018, 2016b, 2013).

### 2.2 Global, linear compensation of cortical thickness for folding effects

As described above, previous studies used FreeSurfer-generated mean curvature (𝐻) to examine the relationship between cortical thickness and curvature and compensated for this correlation by modeling the relationship as 𝑡 = 𝐻 ∗ 𝑘 + 𝑏, where 𝑡 is the uncompensated (i.e., original) cortical thickness, 𝑘 is the regression coefficient, and 𝑏 is the intercept (Glasser et al., 2016; Glasser and Van Essen, 2011; Sigalovsky et al., 2006). The globally compensated thickness was calculated as 𝑡_corr_ = 𝑡 − 𝐻 ∗ 𝑘. Figure 1 illustrates that regressing out mean curvature alone provides only limited compensation for the influence of cortical folding, as measured by mean curvature, on cortical thickness. Fig. 1A displays mean curvature on lateral, medial, and zoomed views of the anterior cingulate cortex on an inflated left hemisphere surface of an individual HCP-YA subject. Fig. 1B shows a map of cortical thickness for the same subject, and the highlighted vertices along gyral ridges (white in Fig. 1A) are consistently thick (red) in Fig. 1B. Fig. 1C shows globally compensated cortical thickness after regressing out mean curvature for this individual. It differs only modestly from the uncompensated thickness in Fig. 1A, whereas it differs dramatically from the group-average cortical thickness after regressing out mean curvature (Fig. 1D), which represents an estimate of the expected pattern if compensation were complete (as residual incompatible folding variability in cortical thickness largely averages out).

**Figure 1:**
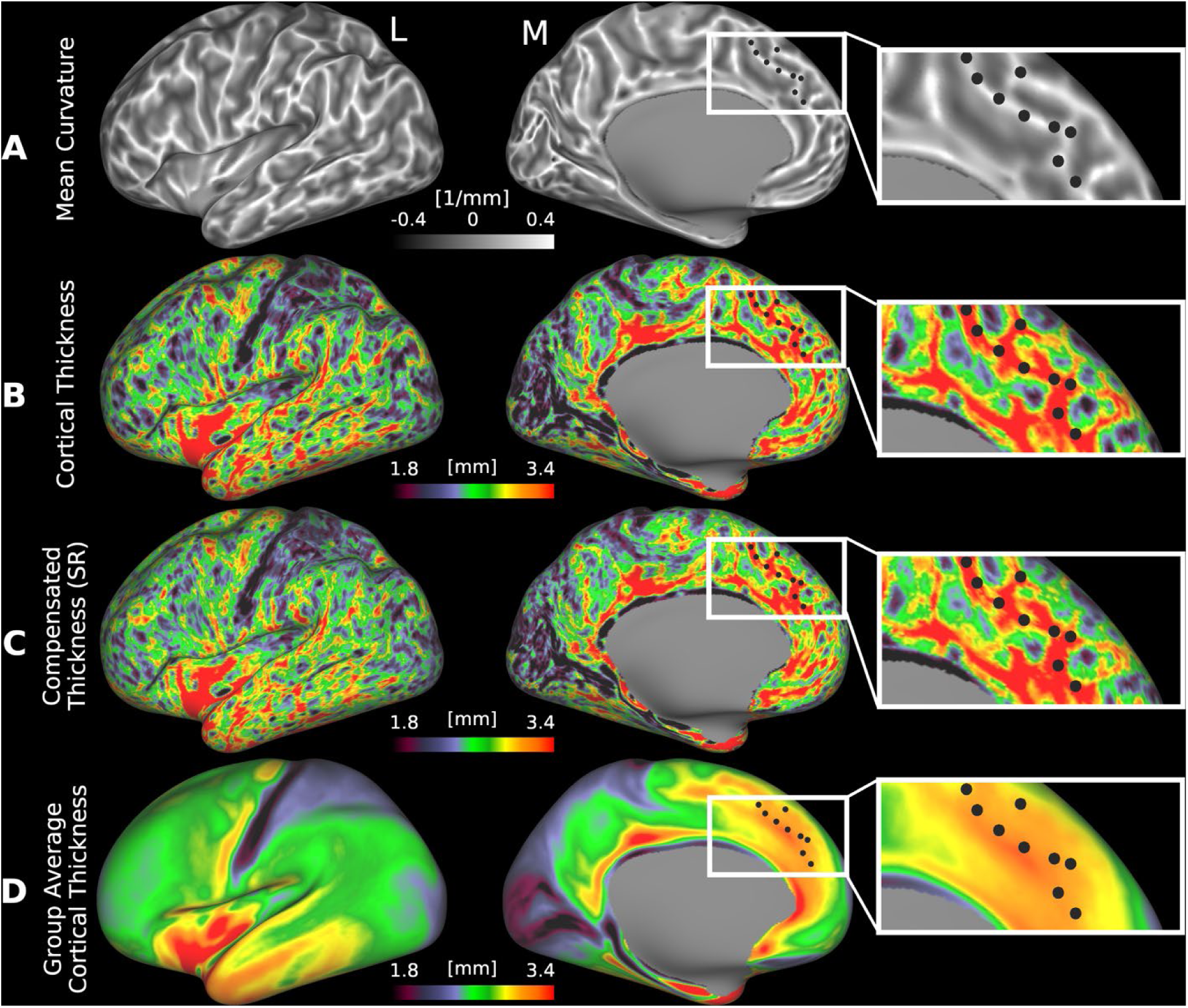
Spatial maps of (A) mean curvature, (B) uncompensated cortical thickness, and (C) globally compensated cortical thickness on the inflated cortical surface of a representative individual from the HCP-YA cohort (case 100408). Insets of the medial prefrontal cortex highlight the spatial correspondence among maps, where regions with high positive curvature (e.g., gyri, bright areas in A) tend to appear thicker (red) in traditional thickness maps (B), an effect slightly attenuated in the globally compensated thickness map (C). (D) shows the group-average map of globally compensated cortical thickness, which illustrates the spatial distribution of cortical thickness after a partial correction of folding effects at the individual level and a marked attenuation of folding-related cortical thickness effects at the group level due to averaging of non-corresponding folding patterns across individuals. The black dots show the corresponding locations in each image. (L: Lateral, M: Medial, SR: Single Regression). Data are available at http://balsa.wustl.edu/mwML7.

### 2.3 Measures of cortical folding

Given that global, linear regression of mean curvature alone is insufficient to fully remove folding effects from cortical thickness, we computed several additional measures of cortical folding. Specifically, we computed five curvature measures that are not collinear with each other: the maximum and minimum principal curvatures (𝑘_1_, 𝑘_2_), Gaussian curvature (𝐾), shape index (𝑆I), and curvedness (𝐶). Importantly, mean curvature (𝐻) is entirely collinear with the maximum and minimum principal curvatures (𝑘_1_, 𝑘_2_) and hence we do not use it in our regression model. Computation of curvature measures for discrete triangular surfaces are explained in detail elsewhere (Demirci and Holland, 2022). In brief, the mean and Gaussian curvatures are computed at each vertex of the midthickness surface (average of white and pial surfaces) using the principles of discrete differential geometry via equations 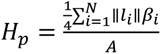 and 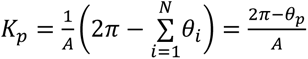 respectively (See Fig. 2A, modified from Demirci and Holland (2022) and Fig. 2B); where 𝑝𝑝 is the vertex in consideration; ‖𝑙_𝑖_‖ is the length of edge 𝑖 connected to vertex 𝑝; 𝛽_𝑖𝑖_ is the dihedral angle, or the angle between normal vectors of the adjacent triangles meeting at the edge 𝑙_𝑖_, (or the acute angle between the two triangular planes); 𝐴𝐴 is the sum of the areas of the adjacent triangles; 𝜃_𝑝_ is the sum of internal angles meeting at vertex 𝑝 (Demirci and Holland, 2022; Mesmoudi et al., 2012). The direction of the unit normal (𝑁_1_, 𝑁_2_ in Fig. 2B) provides the orientation at a given point, so that the mean curvature distinguishes between convex (gyri) and concave (sulci) shapes. Gaussian curvature is positive for both inner concave and outer convex folds (i.e., a cup and a cap), whereas negative values of Gaussian curvature indicate local saddle points, where the surface curves in opposite directions.

**Figure 2:**
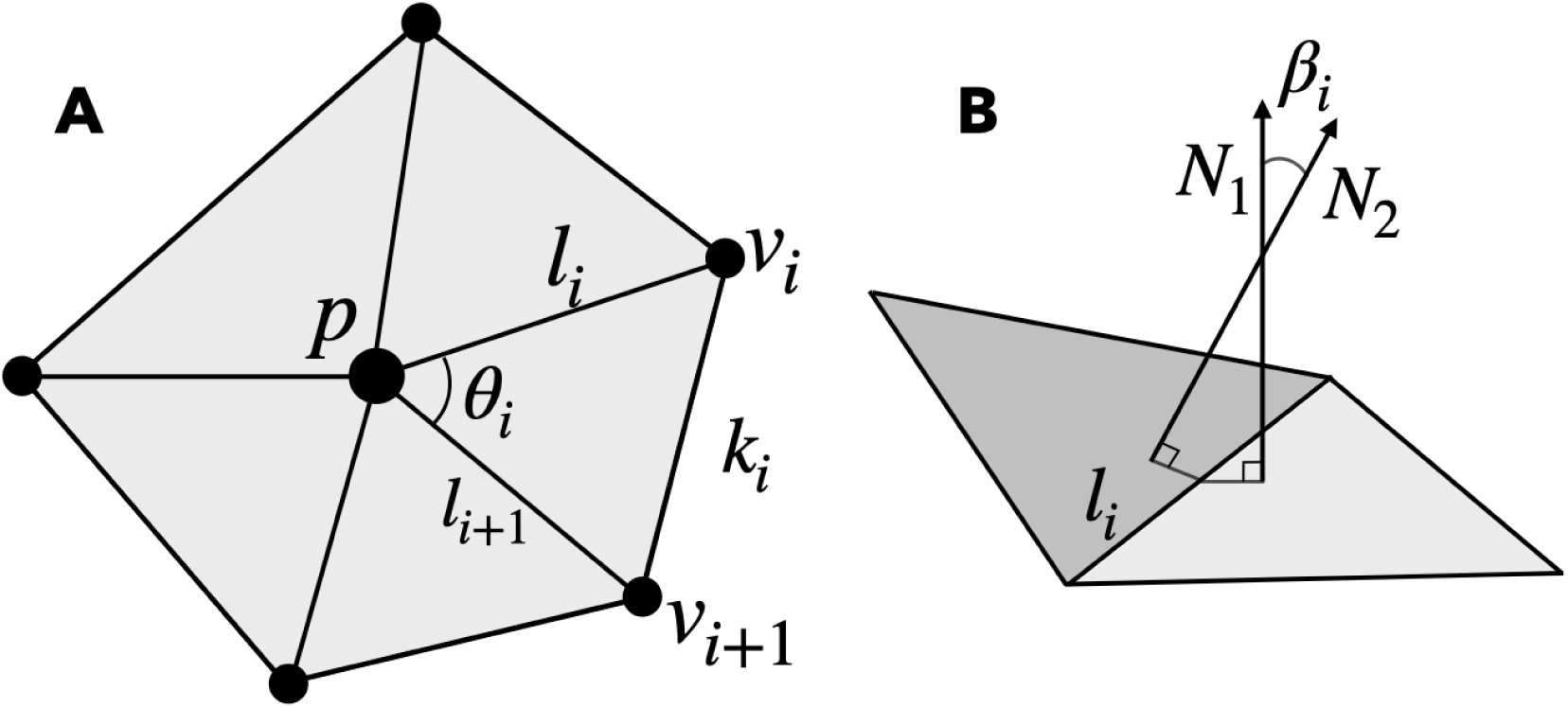
Representative polyhedron of the surface mesh. (A) The center-vertex 𝑝, neighboring vertices 𝑣, edge lengths 𝑙 and 𝑘, and the internal angle 𝜃 are shown (𝑝 is not necessarily co-planar with the neighboring vertices). Modified from Demirci and Holland (2022) (B) The angle 𝛽 between the unit normals (𝑁) of adjacent triangles meeting at edge *l*.

From mean and Gaussian curvatures, the maximum and minimum principal curvatures, 𝑘_1_ and 𝑘_2_, are computed via equations 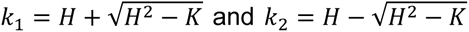, respectively. Then, shape index (𝑆*I*) and curvedness (𝐶) at each vertex are computed as 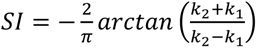 and 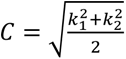, respectively (Koenderink and van Doorn, 1992). The shape index is a dimensionless scalar measure, ranging from −1 to 1, mapping local surface shapes continuously based on principal curvatures, providing a continuous way to classify local surface features such as peaks (spherical cap or dome), ridges, saddles, ruts, and valleys (spherical cup or trough). The dimensionless shape index allows comparison of cortical folding patterns across different-sized brains, for example, during development, aging, or across species. Curvedness on the other hand is dimensional, always positive, and specifies the magnitude of surface curvature. It is inversely proportional to the size of the object (1⁄𝑚𝑚) and vanishes only at planar points (Koenderink and van Doorn, 1992). Curvedness is higher for highly folded regions such as sharp outer ridges and deep inner valleys, and low for gently curved or nearly flat regions.

The five curvature measures used in the computation of curvature-compensated cortical thickness are shown in Fig. 3. Maximum principal curvature (𝑘_1_) is always greater than the minimum principal curvature (𝑘_2_) at each point. High values of |𝑘_1_| correspond to prominent gyral crowns (Fig. 3A), whereas high values of |𝑘_2_| correspond to sulcal fundi (Fig. 3B). As 𝑘_1_ ≥ 𝑘_2_, maximum principal curvature highlights outward bends, which are mostly found on gyri (as the bending in the other direction is minor due to its cylindrical shape), and as |𝑘_2_| ≥ 𝑘_1_ the minimum principal curvature highlights inward bends, which are mostly found in sulci. However, it is important to note that both inward and outward bends can be found at any location throughout the cortex (e.g., the insula has its own gyri and sulci). Given that cortical thickness varies systematically across gyri and sulci—generally thicker at gyral crowns and thinner at sulcal fundi—the relationship between folding and thickness is expected to be directionally opposite for these two measures. (*K*) can occur in sulcal and gyral regions, reflecting different local surface geometries. Specifically, *K* > 0 indicates either convex (cap/dome-shaped) or concave (cup/trough-shaped) regions, whereas *K* < 0 reflects saddle-like configurations, typically found at transitions between convex and concave areas (Fig. 3C). The shape index, on the other hand, (*SI*) captures these distinctions across three primary geometric types: convex (*SI* > 0), concave (*SI* < 0), and saddle-shaped (*SI* ≈ 0) regions (Fig. 3D). Finally, curvedness (*C*) which is always positive, quantifies the overall magnitude of surface bending and increases with sharper local curvature (Fig. 3E).

**Figure 3:**
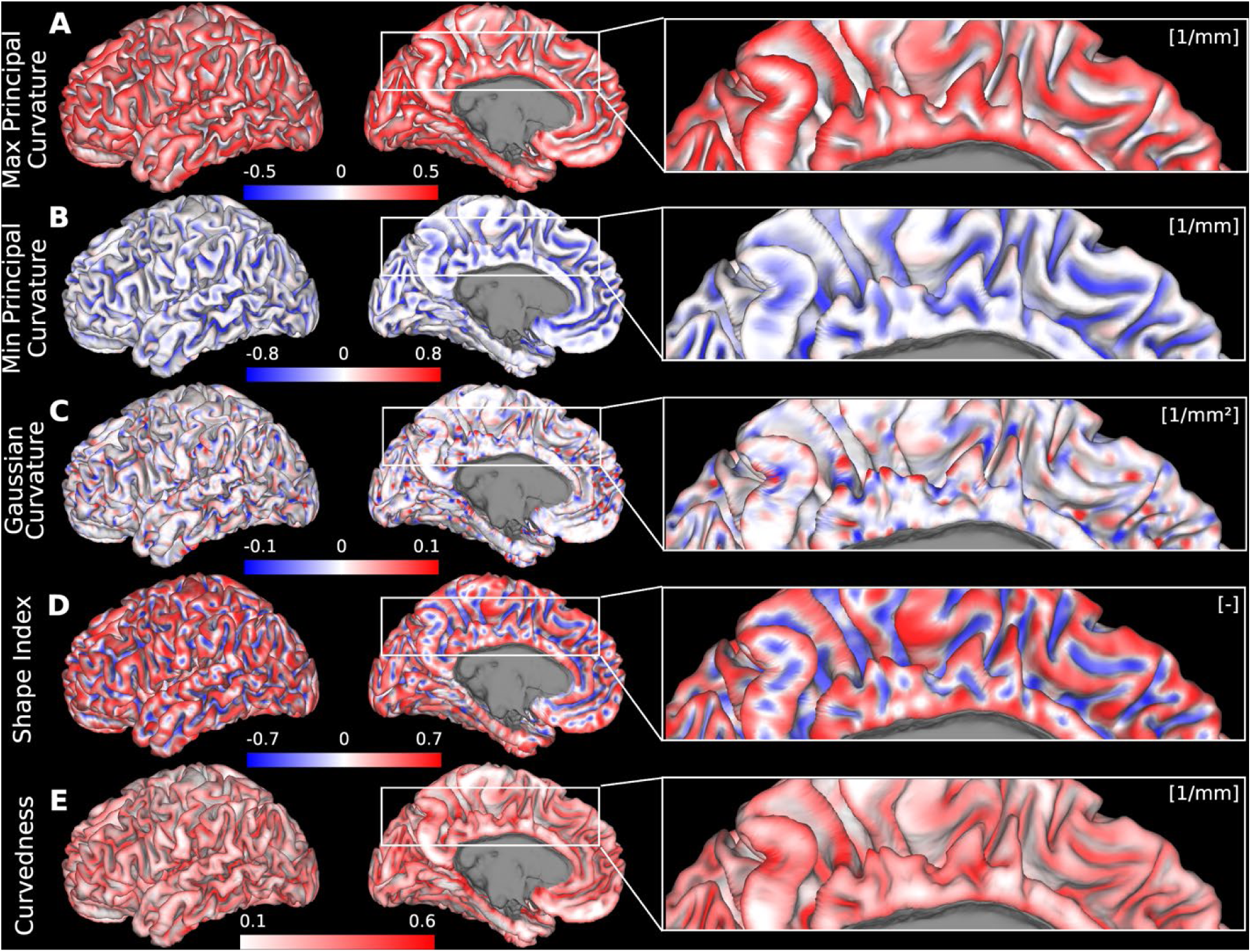
The five curvature measures used in the multiple regression model. (A) Maximum principal curvature, (B) minimum principal curvature, (C) Gaussian curvature, (D) shape index, and (E) curvedness. Spatial maps of each curvature are displayed on the midthickness surface of a representative individual (case 100408) from the HCP-YA cohort. Data are available at http://balsa.wustl.edu/Vlg57.

### 2.4 Regularization of Folding Measures

The native FreeSurfer meshes are highly irregular in vertex spacing and density because they are derived from the tessellation of the segmented voxels. Instead of using the native Freesurfer meshes, we computed the folding measures after first resampling the midthickness surface (average of the coordinates of the grey/white and pial surfaces (Van Essen et al., 2012)) to a regular 164k vertex standard mesh (without performing a surface registration), using the standard barycentric resampling algorithm available in Connectome Workbench (Fig. 4A). This process preserves the shape of the midthickness surface mesh while ensuring that roughly similar surface areas are associated with each vertex and nearly all vertices have six neighbors. Next, the midthickness surface mesh was slightly spatially smoothed by applying a geodesic Gaussian kernel of a given FWHM. This smoothing reduces the effects of noise from the structural images and other slight irregularities in the surface placement (Fig. 4B). The smoothing operation is similar to Taubin smoothing (Taubin, 1995), where surface shrinkage is avoided and fine surface details are preserved. These steps further enhance the accuracy and robustness of vertex-wise curvature calculations, which are highly dependent on the quality of the triangular mesh. The numerical stability of these calculations relies on the homogeneity of the triangular aspect ratio, size, and density. Highly skewed or elongated triangles can introduce numerical artifacts in curvature estimations, and uneven vertex distribution can result in local inaccuracies. Next the curvature data are computed as described above and smoothed on the surface using a geodesic Gaussian kernel of a given FWHM to reduce noise and mitigate the impact of extreme outliers (such as isolated spikes), ensuring their influence is minimized while maintaining spatial consistency without distorting the true distribution (Fig. 4C).

**Figure 4:**
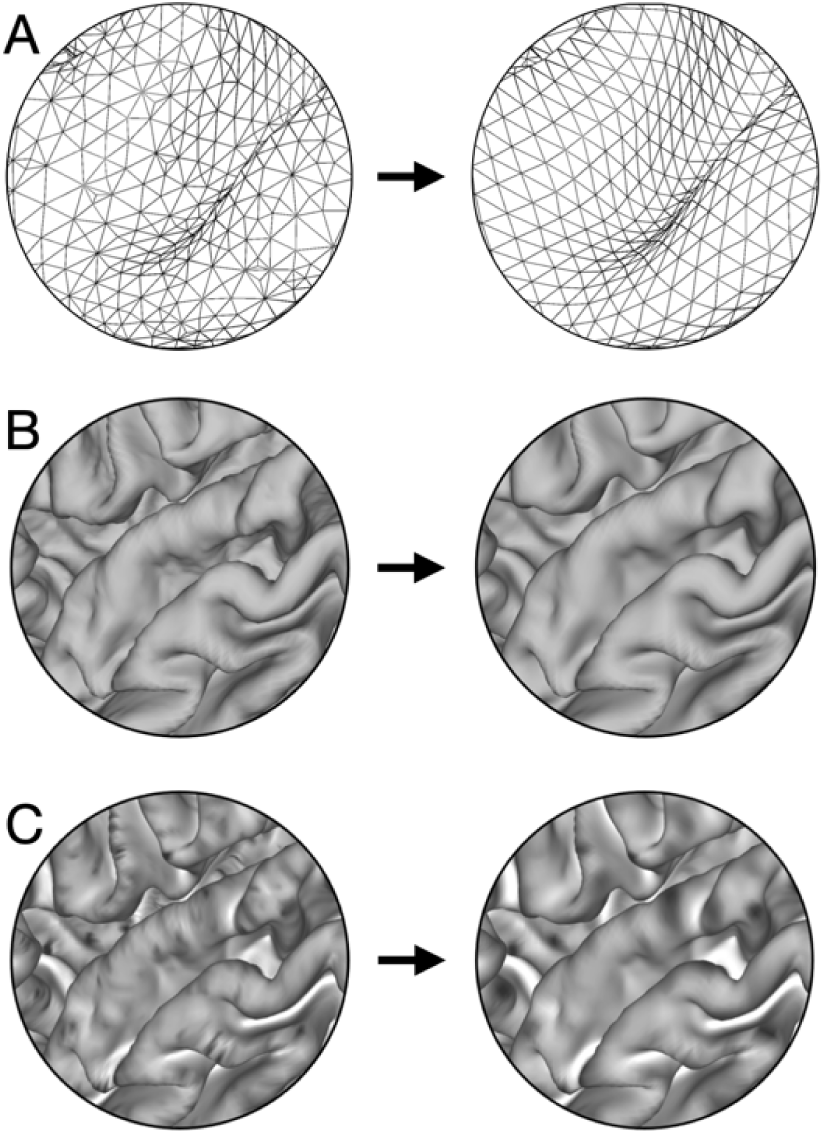
Regularization of cortical folding measures. (A) Resampling of the native FreeSurfer midthickness surface to a standardized 164k-vertex mesh. (B) Surface-based smoothing of the midthickness geometry. (C) Smoothing of curvature measures, illustrated here with the maximum principal curvature.

### 2.5 Local nonlinear multiple regression model for compensating cortical thickness for folding effects

The above five measures of folding and their squared terms are demeaned and included in the multiple regression model, solving the equation: 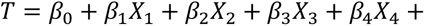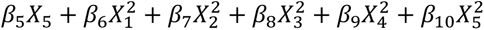 for unknown coefficients 𝛽_𝑖_ . 𝑇 is the original (uncompensated) cortical thickness, 𝑋_1_, 𝑋_2_, 𝑋_3_, 𝑋_4_, 𝑋_5_ are predictors (i.e., curvatures), 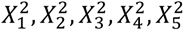 are the squared terms of each predictor so as to include both linear and quadratic effects in the regression model (Power et al., 2014). The polynomial order was determined by evaluating the relationship between cortical thickness and curvature descriptors. As shown by the regression coefficient maps (Supplementary Figure 2), second-order terms were close to zero, indicating that the relationship is predominantly linear and it would not be productive to include higher-order terms in the regression equation. The multiple regression is defined uniquely for each vertex and is weighted based on a Gaussian of a given FWHM centered at that vertex (Fig. 5A). For each region centered at the vertex, we ensured by visual inspection that in general at least one outward and one inward fold were included, such that our lowest search range was 3 mm FWHM for the regional patch size (See Supplementary Figure 1). This local approach to the multiple regression enables the compensation model to flexibly fit the folding effects based on regional variation in the uncompensated cortical thickness, avoiding the underfitting seen with the global linear regression approach. Finally, the curvature-compensated thickness is calculated by solving the equation 𝑇_corr_ = 𝑇*_orig_* − 𝛽_1:5_𝐹 − 𝛽_6:10_𝐹^2^, where 𝛽_1:5_ are the linear coefficients, 𝛽_6:10_ are quadratic coefficients, and 𝐹𝐹 are the curvature measures.

**Figure 5:**
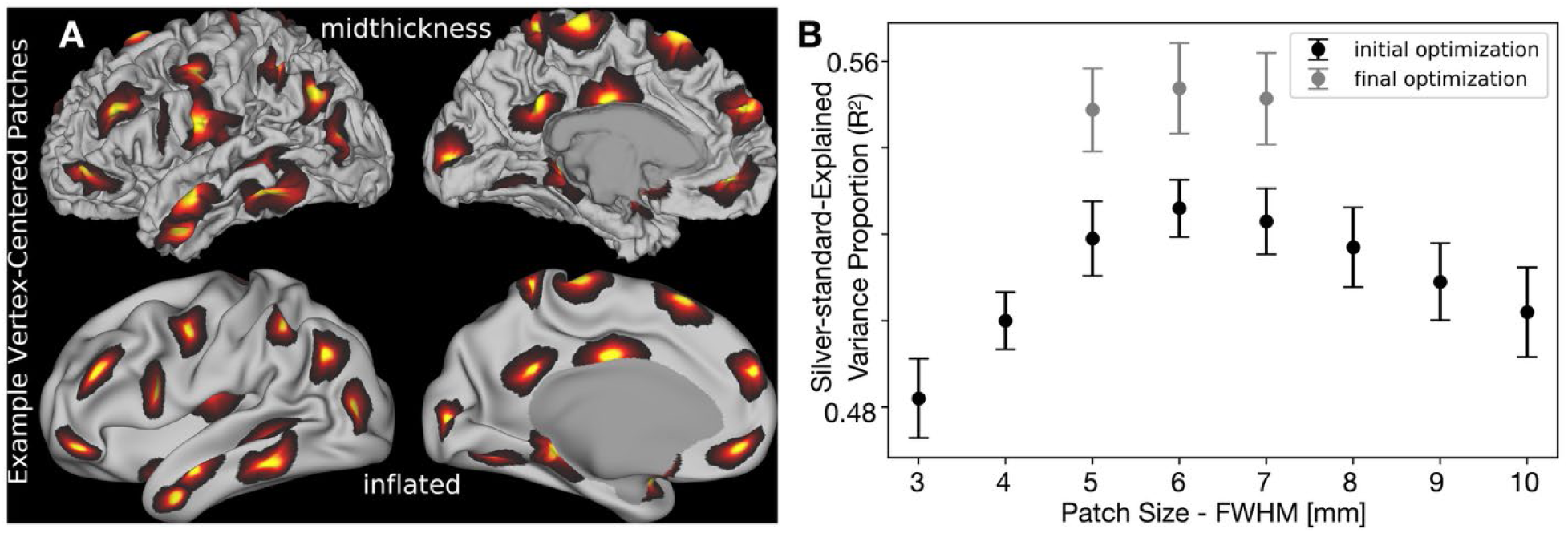
Optimization of patch size for local nonlinear folding compensation of cortical thickness. (A) Example vertex-centered patches (6 mm FWHM) displayed on the midthickness and inflated surfaces of a representative individual from the HCP-YA cohort. Medial wall vertices are masked and excluded from all our analyses. (B) Patch size optimization using a golden-section search algorithm. The initial optimization targets the group-average cortical thickness map following global linear regression-based folding compensation. Final optimization uses the group-average map generated from local nonlinear regression-based folding compensation, initialized with the parameters from the first stage. Error bars represent the variability across the 10 tested individuals. Data are available at http://balsa.wustl.edu/z0vBk.

### 2.6 Optimization of free parameters

Above we defined three free parameters for the curvature measures and the local multiple regression: Gaussian FWHMs for surface smoothing, curvature smoothing, and the vertex-centered weighted local regression. Our objective was to find a consistent set of parameters across individuals that best enable us to estimate what the cortical thickness would have been in the absence of cortical folding. In humans, the high interindividual variability in cortical folding—particularly after areal feature-based alignment such as MSMAll— allows the use of the group-average cortical thickness as a proxy for folding-compensated cortical thickness. Averaging across individuals naturally attenuates the spatially inconsistent effects of cortical folding on cortical thickness. While the global linear regression method provides only a partial compensation at the individual level, it further reduces folding-related effects in the group-average cortical thickness map. Thus, we take the group-average global linear regression folding- compensated cortical thickness map (Fig. 1D) as a *silver standard* and an optimization target for the individual local multiple regression compensation algorithm. We then use the golden-section search algorithm on ten individual HCP-YA participants to explore the parameter space of these three parameters. The optimization criterion for the golden-section search was the proportion of variance explained (𝑅^2^) in the individual folding-compensated map by the silver standard group-average map. The initial search interval for surface and curvature smoothing was determined based on prior empirical knowledge: [0, 3] mm FWHM. Specifically, we sought to avoid over-smoothing by visually inspecting surfaces and curvature maps, as illustrated in Fig. 4. For ROI patch size [3, 10] mm FWHM range was selected; the minimum radius was set to 3 mm to ensure inclusion of at least one inward and one outward fold (See Supplementary Figure 1) and the maximum search radius was set to 10 mm after trial and error, which balanced capturing local geometry with avoiding unnecessary computational burden. The optimization followed a sequential order: 1) patch size, 2) surface smoothing, and 3) metric (i.e., curvature) smoothing. The iteration stopping criterion for the golden-section search optimization was set at 0.1; when the difference between the upper and lower parameter bounds fell below 0.1, the iteration stopped. This value was determined empirically through trial and error to provide sufficient optimization precision and sensitivity without excessive computational cost.

The final optimized parameters are 2.14 mm FWHM for surface smoothing, 2.52 mm FWHM for curvature smoothing, and 6 mm FWHM for the patch size. These values provided the highest 𝑅^2^across the tested individuals with no visible artifacts in the folding-compensated cortical thickness map. Fig. 5B illustrates the optimization function for the Gaussian FWHM of the vertex-centered weighted local regression for the ten individuals. After computing the optimized cortical thickness regression for all 1071 HCP-YA individuals, we re-ran the golden section search using the group-average local multiple regression compensated thickness map on the same 10 individuals and found similar optimal parameters.

### 2.7 Data and Code Availability

The local multiple regression-based folding-compensated cortical thickness pipeline is now part of the HCP Pipelines (Glasser et al., 2013) on GitHub (https://github.com/Washington-University/HCPpipelines) and is now a default output of new runs of the HCP Structural Preprocessing pipelines (specifically the PostFreeSurfer pipeline). The pipeline can also be run on existing HCP Pipelines outputs using the global/scripts/CorrThick.sh pipeline module. An example script is provided at Examples/Scripts/CorrThickPipelineBatch.sh. In the HCP Pipelines’ outputs and HCP data releases, the global linear folding-compensated cortical thickness file contains the tag *corrThickness*. Our new measure of local, nonlinear, folding- compensated cortical thickness file contains the tag *MRcorrThickness*. HCP installation instructions and dependencies are available within the HCP Pipelines GitHub repository documentation, or the HCP Pipelines can be run within the QuNex container (Ji et al., 2023). The data shown in this publication are available as a BALSA study at https://balsa.wustl.edu/study/Dlr9k and the raw HCP data are in ConnectomeDB powered by BALSA.

### 2.8 Computation details

The folding-compensated cortical thickness pipeline takes ∼30 minutes per subject on a Linux system with a 3.1 GHz, 14-core CPU (∼90 minutes on a single core) and requires ∼3 GB of RAM.

## Results

### 3.1 Individual folding-compensated cortical thickness patterns

Figure 6 presents folding as quantified by the shape index (Fig. 6A), cortical thickness (Fig. 6B), globally compensated cortical thickness using a linear regression approach (Fig. 6C), and locally compensated cortical thickness using a nonlinear multiple regression model (Fig. 6D) on an inflated cortical surface from the same HCP-YA individual as in Fig. 1. As shown in Figures 6A and 6B, uncompensated cortical thickness patterns closely follow folding patterns, with negatively curved regions (sulcal fundi) exhibiting markedly lower thickness than positively curved regions (gyral crowns). Applying the global linear compensation (Fig. 6C) reduces the strength of this relationship somewhat but does not fully eliminate it. In contrast, after the local nonlinear compensation (Fig. 6D), the previously strong association between cortical thickness and folding is no longer visually apparent. Instead, the compensated thickness maps appear more spatially homogeneous across gyri and sulci, suggesting that the local, nonlinear model more effectively accounts for folding-related variability in cortical thickness. Furthermore, the standard deviation of local nonlinear compensated thickness, CT(MR) (𝜎 = 0.42 𝑚𝑚) is lower than that for cortical thickness, CT (𝜎 = 0.57 𝑚𝑚) or global linear regression compensated thickness, CT(SR) (𝜎 = 0.54 𝑚𝑚), for an individual from the HCP-YA cohort (Fig. 6E) providing further evidence that the nonlinear local correction approach effectively reduces folding-driven variability across the cerebral cortex. This reduction in variability suggests that local nonlinear compensation accounts for region-specific effects that are not fully captured by a global linear model, resulting in a more biologically meaningful representation of individual intrinsic cortical thickness patterns. Much of the remaining high-spatial-frequency variation in the multiple nonlinear, local regression-based compensation thickness estimate is related to sub-mm errors in surface placement.

**Figure 6:**
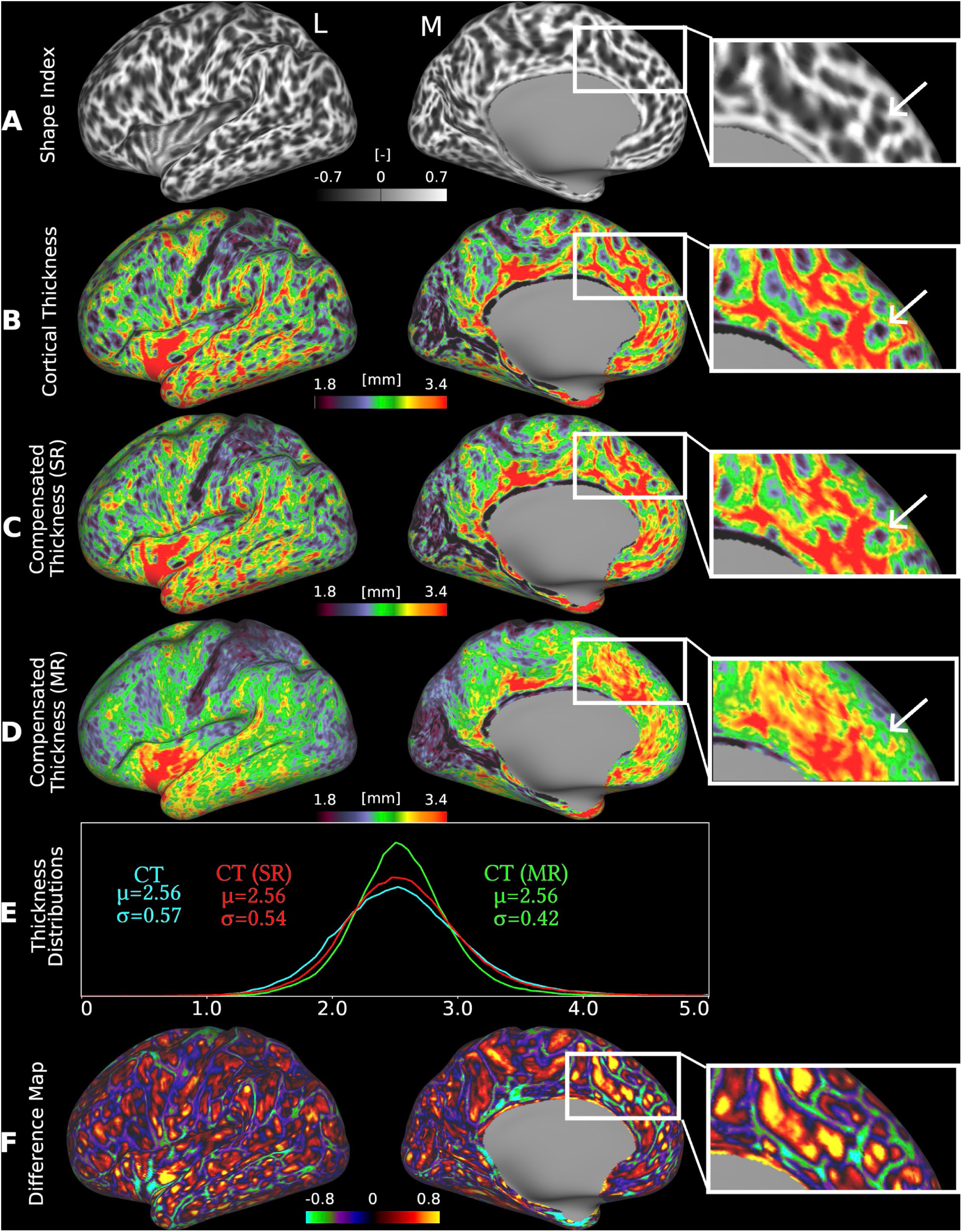
Folding compensation of cortical thickness. (A) shape index, (B) cortical thickness, (C) cortical thickness after global, linear regression-based folding compensation (SR: Single Regression), and (D) cortical thickness after local, nonlinear regression-based folding compensation (MR: Multiple Regression) are shown on the inflated surface of a representative individual from the HCP-YA cohort (case 100408). Zoomed insets in A, B, C, and D highlight the pronounced differences between global and local compensation approaches, with the local nonlinear method more effectively reducing folding-related bias than the global linear method (white arrows). (E) Comparison of thickness distributions reveals reduced intra-individual variability following local nonlinear folding compensation. (F) Spatial difference map between uncompensated and locally compensated cortical thickness, illustrating the effect of folding compensation. (L: Lateral, M: Medial). Data are available at http://balsa.wustl.edu/XZgKz and http://balsa.wustl.edu/8mX52.

### 3.2 Uncompensated and compensated cortical thickness difference map

Figure 6F illustrates the difference between uncompensated and local nonlinear compensated cortical thicknesses. The first-order effects of the map are that sulci get thicker with compensation and gyri get thinner, and that sulci and gyri change by similar amounts. The difference map also reveals a second-order effect in which changes are more pronounced in cortical regions that are thick on average (e.g., insular cortex, anterior temporal cortex, anterior cingulate cortex and medial prefrontal cortex), whereas changes are more modest in cortical regions that are thin on average (e.g., visual cortex, superior parietal cortex), suggesting that reduced compensation is needed in these areas. These findings underscore the limited effectiveness of global compensation and highlight the need for the localized (i.e., regional) correction strategy employed here.

### 3.3 Folding-compensated and uncompensated cortical thickness patterns for 1071 healthy young adults

We examined the interactions between cortical shape, sulcal depth, and cortical thickness in individuals from the HCP-YA cohort. Cortical thickness is strongly correlated with shape and varies with depth, as deeper sulcal fundi tend to have thinner cortices than outer gyral crowns. Notably, the correlation between shape and thickness for a given sulcal depth, and the correlation between depth and thickness for a given shape, are consistent (Demirci and Holland, 2022). To test this relationship on uncompensated and folding-compensated thickness measures, we analyzed all 1071 HCP-YA individuals. Unlike the previous work, we used Freesurfer-generated *sulc* as a proxy for *sulcal depth*. The *sulc* metric reflects the relative displacement of each vertex from an estimated mid-surface between gyri and sulci, providing a measure of the depth and height of cortical folds. It serves as a proxy of linear distance but is dimensionless, unlike a true sulcal depth measure. Using the shape index and sulcal depth ranges indicated in Fig. 7, we computed each individual’s mean cortical thickness and then averaged it across the cohort. If no vertices were available in a given range for an individual (e.g., sharply convex points in deep sulcal bends), that individual was excluded from the group average. Fig. 7 reveals a systematic variation of cortical thickness with both shape and depth, similar to the prior work (Demirci and Holland, 2022). Specifically, cortical thickness increases gradually from convex to concave regions and from the bottom of sulci to the crest of gyri (yellow to blue gradient going from top left to bottom right in Fig. 7A). A similar analysis using the global, linear folding-compensated cortical thickness shows a more homogeneous cortical thickness patterning, but the relationship between the three measures is still discernible (Fig. 7B). In contrast, when considering local, nonlinear folding-compensated cortical thickness, this systematic patterning disappears, and no clear relationship with shape or depth is observed. Instead, local, nonlinear folding-compensated cortical thickness appears uniform and spatially homogeneous across the whole cortex (Fig. 7C).

**Figure 7:**
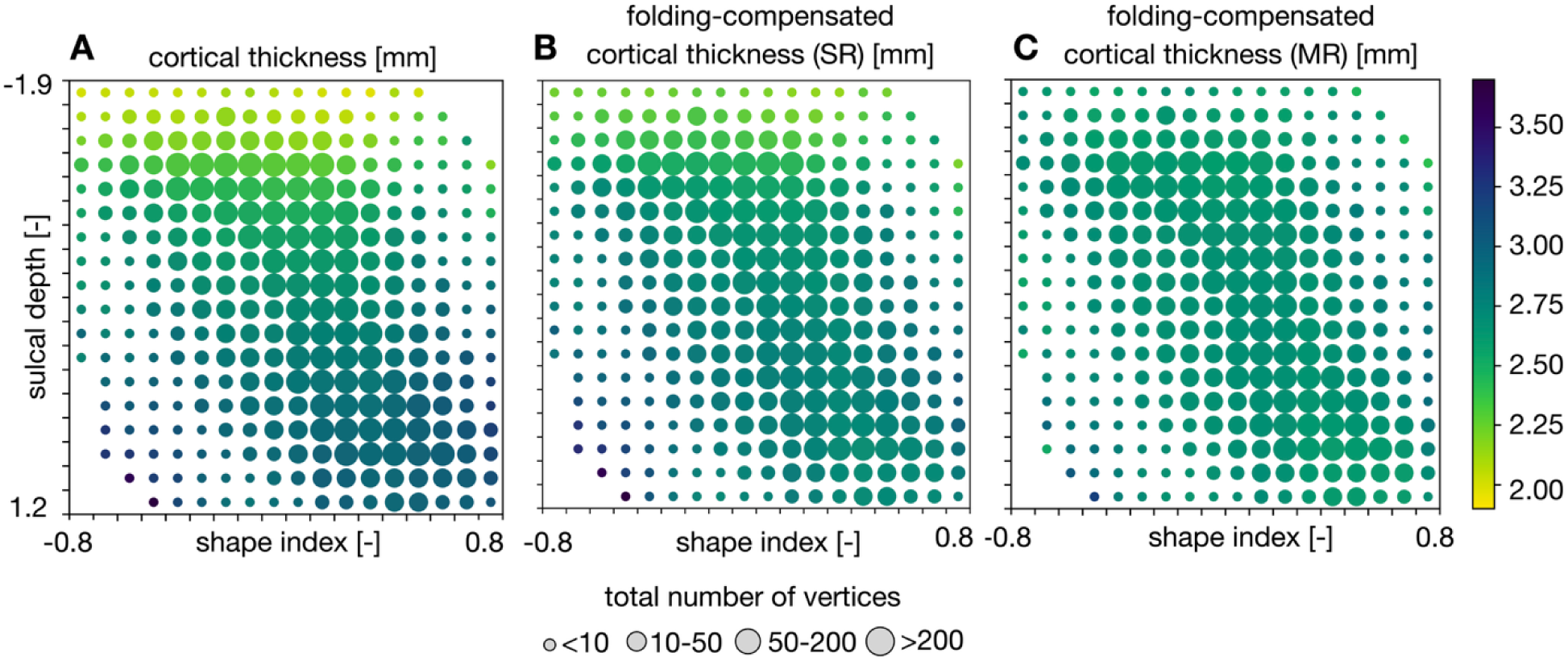
Folding-compensated and uncompensated cortical thickness patterns across all HCP-YA individuals, shown with respect to shape and depth (vertex location) features. (A) Uncompensated cortical thickness. (B) Cortical thickness after global linear folding compensation using single regression (SR). (C) Cortical thickness after local nonlinear folding compensation using multiple regression (MR). For each range of sulcal depth and shape index, we computed each individual’s mean cortical thickness and averaged it across the cohort. Note that, if no vertices are available within the given range, that individual is excluded from the group average. The local nonlinear method yields more uniform thickness estimates by more effectively accounting for individual variability in cortical folding.

### 3.4 Group-average folding-compensated cortical thickness patterns for healthy young and aging adults

We computed the group-average folding-compensated cortical thickness patterns for the HCP-YA and HCA cohorts, as shown in Figures 8A and 8B, respectively, where folding-compensated cortical thickness here refers to the locally compensated cortical thickness obtained using multiple regression. In both groups, the spatial distribution of folding-compensated cortical thickness appears more uniform compared to patterns of uncompensated cortical thickness, particularly in the somatosensory, medial prefrontal, and lateral temporal cortices. This suggests a reduction in the influence of cortical folding on group-level patterns, while preserving biologically meaningful regional specializations, i.e., thicker motor and thinner somatosensory cortices, consistent with known cytoarchitectonic differences.

**Figure 8:**
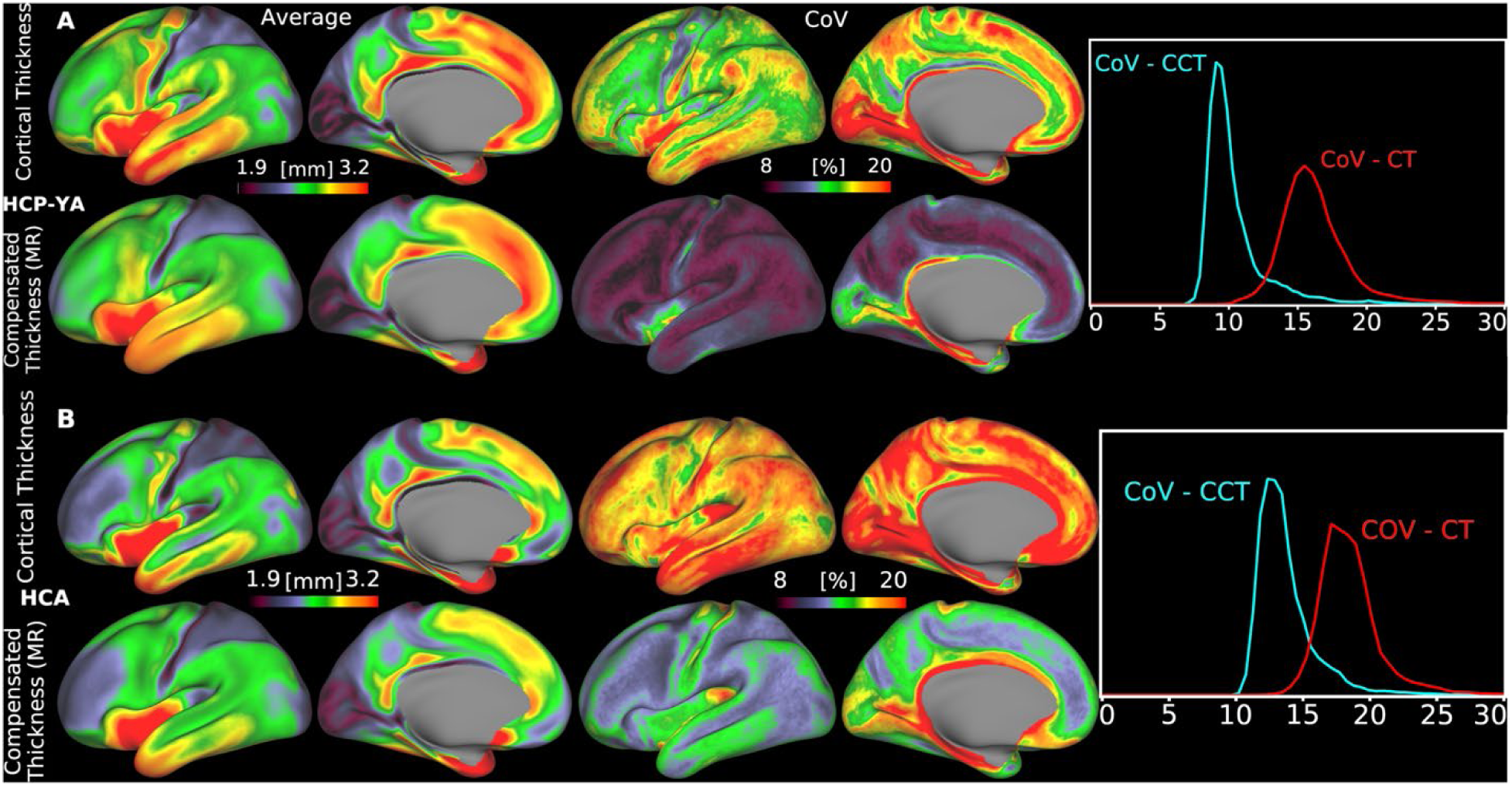
Group-average cortical thickness patterns before and after folding compensation, along with their corresponding coefficients of variation (CoV) across individuals, shown for (A) the HCP-YA cohort and (B) the HCA cohort. Folding-compensation of cortical thickness reduces inter-individual variability, as reflected by lower CoV values (MR: Multiple Regression). Data are available at http://balsa.wustl.edu/NP5Bn and http://balsa.wustl.edu/G70B1.

To quantify the effect on individual variability, we computed the coefficient of variation (CoV), defined as the standard deviation normalized by the mean, at each cortical vertex for both groups. Across the cortex in both HCP-YA, Fig. 8A and HCA, Fig. 8B, CoV is markedly lower for folding- compensated cortical thickness (*mean* = 10%; cyan curve in Fig. 8A and *mean* = 13%; cyan curve in Fig. 8B) compared to uncompensated thickness (*mean* = 15%; red curve in Fig. 8A and *mean* = 17%; red curve in Fig. 8B), indicating a marked reduction of inter-individual differences in cortical thickness that are caused by inter-individual folding variability. However, CoV remains high in specific regions, particularly the insular, visual, and somatosensory cortices. These regions generally have greater inaccuracies in surface placement due to having thin, heavily myelinated cortex or thin white matter overlying the claustrum in insular cortex. The greater CoV in HCA vs HCP-YA subjects in both compensated and uncompensated thickness may in part reflect technical factors such as the slightly coarser voxel size (0.8 mm vs 0.7 mm isotropic) and the greater age range studied by HCA.

Finally, age-related cortical atrophy is evident in both maps, with a very similar pattern (spatial correlation 𝑟 = 0.92; Fig. 9). The most pronounced reductions are in medial prefrontal cortex, which is consistent with established patterns of age-related neurodegeneration (Hurtz et al., 2014; Salat et al., 2004) and shows that folding compensation of cortical thickness does not substantially alter a key neurobiological finding of interest.

**Figure 9:**
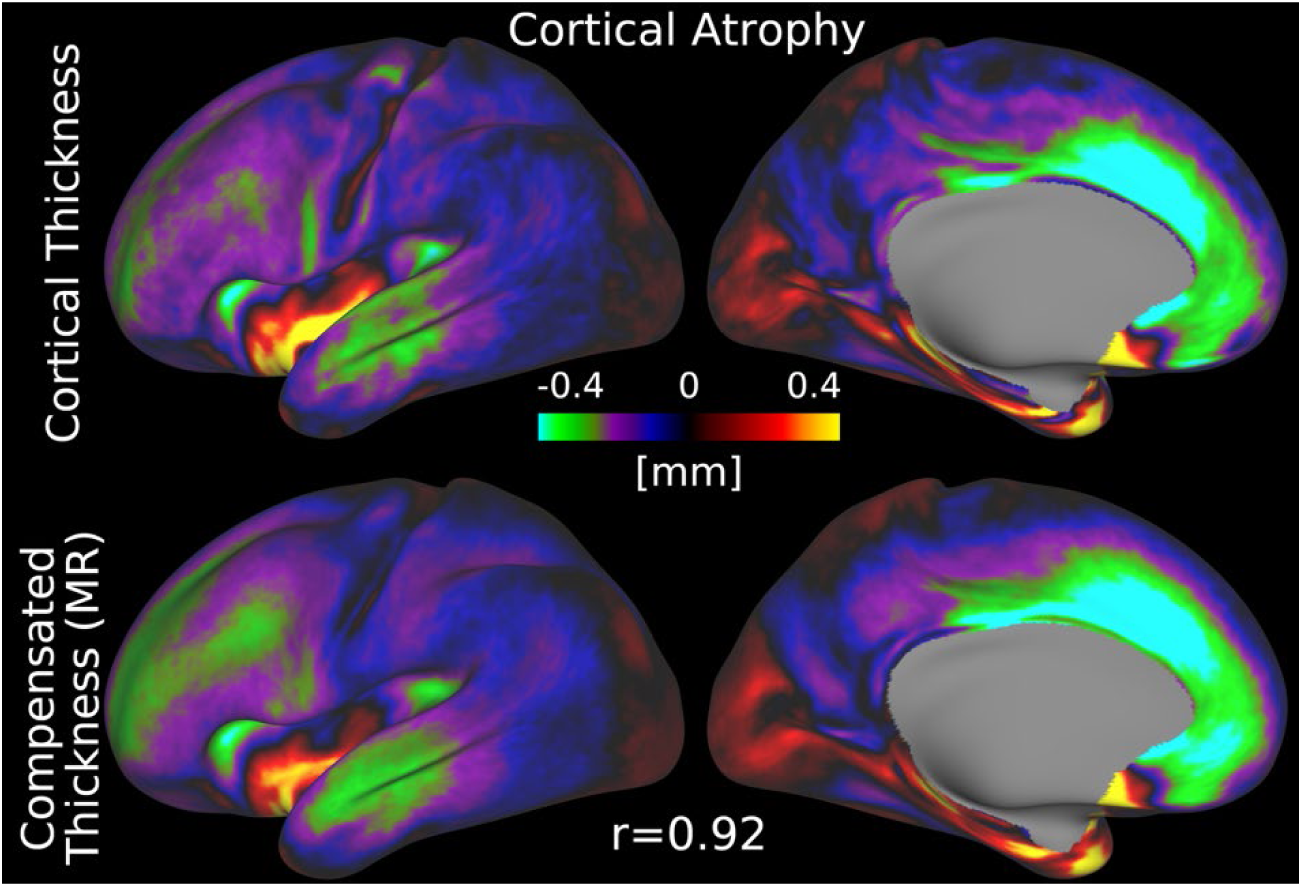
Age-related cortical atrophy based on uncompensated and folding-compensated cortical thickness. Spatial maps depict the difference between HCA and HCP-YA group averages for both measures. Both maps exhibit a highly similar spatial pattern, with a strong correlation (𝑟 = 0.92) (MR: Multiple Regression). Data are available at http://balsa.wustl.edu/L38pz.

### 3.5 Parcellated folding-compensated cortical thickness for healthy young and aging adults

We further examined the differences in spatial patterns between parcellated cortical thickness and folding-compensated cortical thickness in HCP-YA (Fig. 10) and HCA (Fig. 11), where folding- compensated cortical thickness here refers to the locally compensated cortical thickness obtained using multiple regression. For both cohorts, we computed the mean of both the mean thickness and the standard deviation in cortical thickness within each cortical area across individuals. Overall, the spatial distribution of mean areal cortical thickness is similar for the two measures and is consistent across cohorts (Figs. 10A & 11A). Importantly, the mean areal standard deviation is markedly lower for folding-compensated cortical thickness in both groups. In HCP-YA, the mean areal standard deviation decreases from 0.38 mm (uncompensated) to 0.22 mm (compensated) (Fig. 10B), and in HCA, from 0.40 mm to 0.25 mm (Fig. 11B). This indicates that the folding compensation makes cortical thickness more homogeneous within cortical areas that span both gyral and sulcal regions. We also compared the left and right hemispheres in terms of areal mean and variability in both groups (Fig. 12) and observed a strong hemispheric symmetry between the left and right hemispheres for both compensated and uncompensated cortical thickness. The areas with highest and lowest mean and standard deviation are the same for both measures and groups, as shown in Fig. 12.

**Figure 10:**
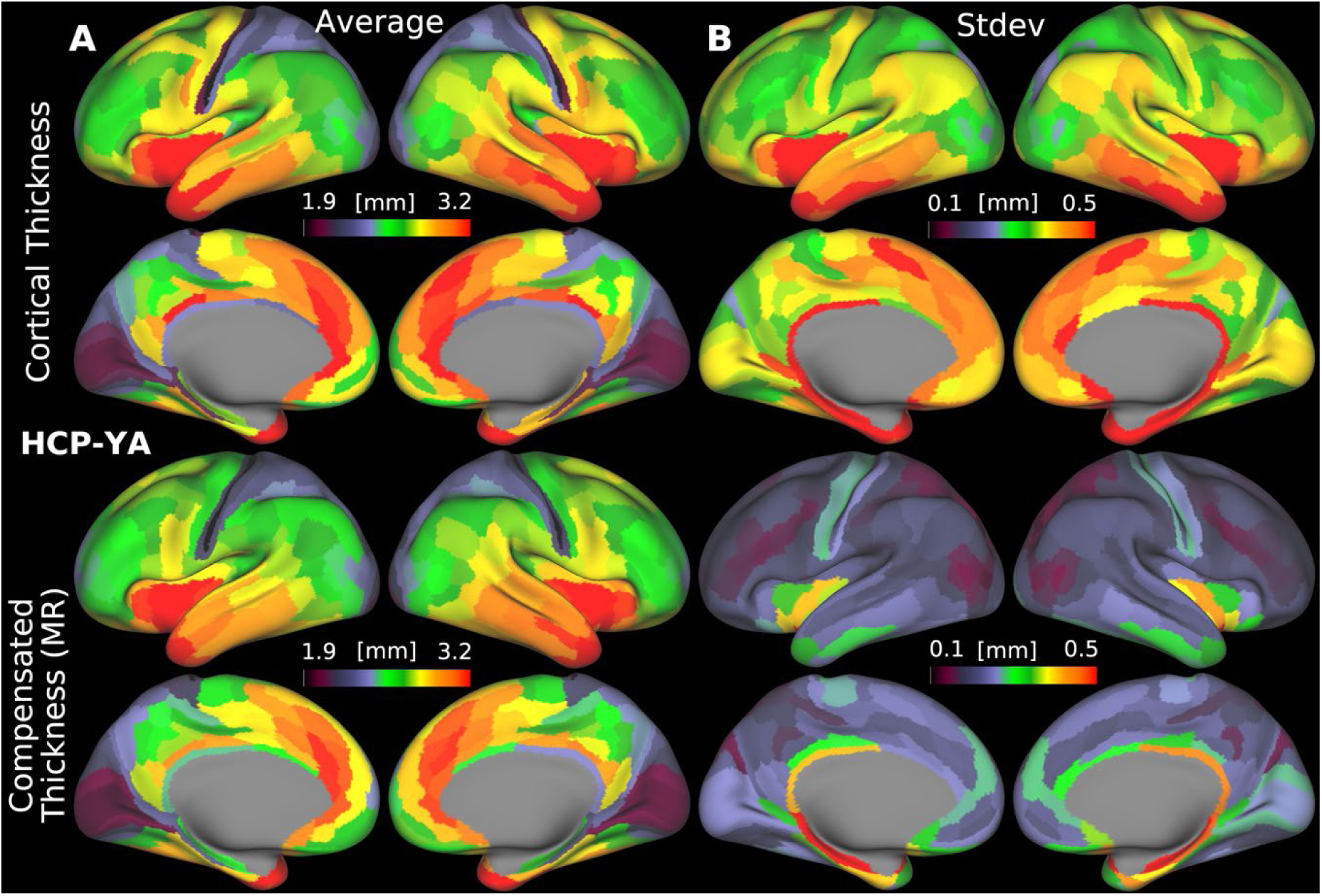
Parcellated uncompensated and folding-compensated cortical thickness for the HCP-YA cohort. Group areal mean (A) and group mean areal standard deviation (B) are shown (MR: Multiple Regression). Data are available at http://balsa.wustl.edu/pXMr7.

**Figure 11:**
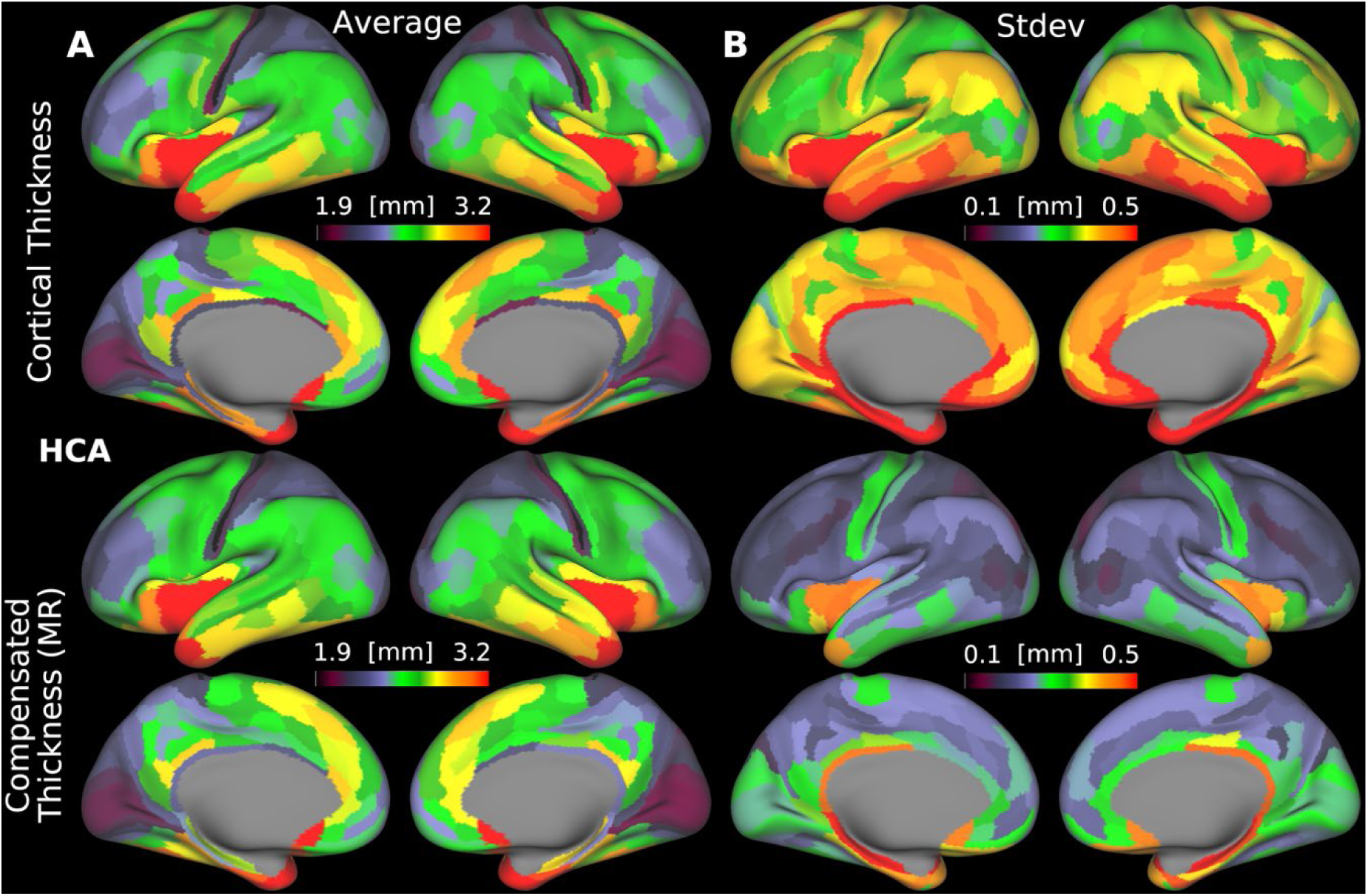
Parcellated uncompensated and folding-compensated cortical thickness for the HCA cohort. Group areal mean (A) and group mean areal standard deviation (B) are shown (MR: Multiple Regression). Data are available at http://balsa.wustl.edu/9648D.

**Figure 12:**
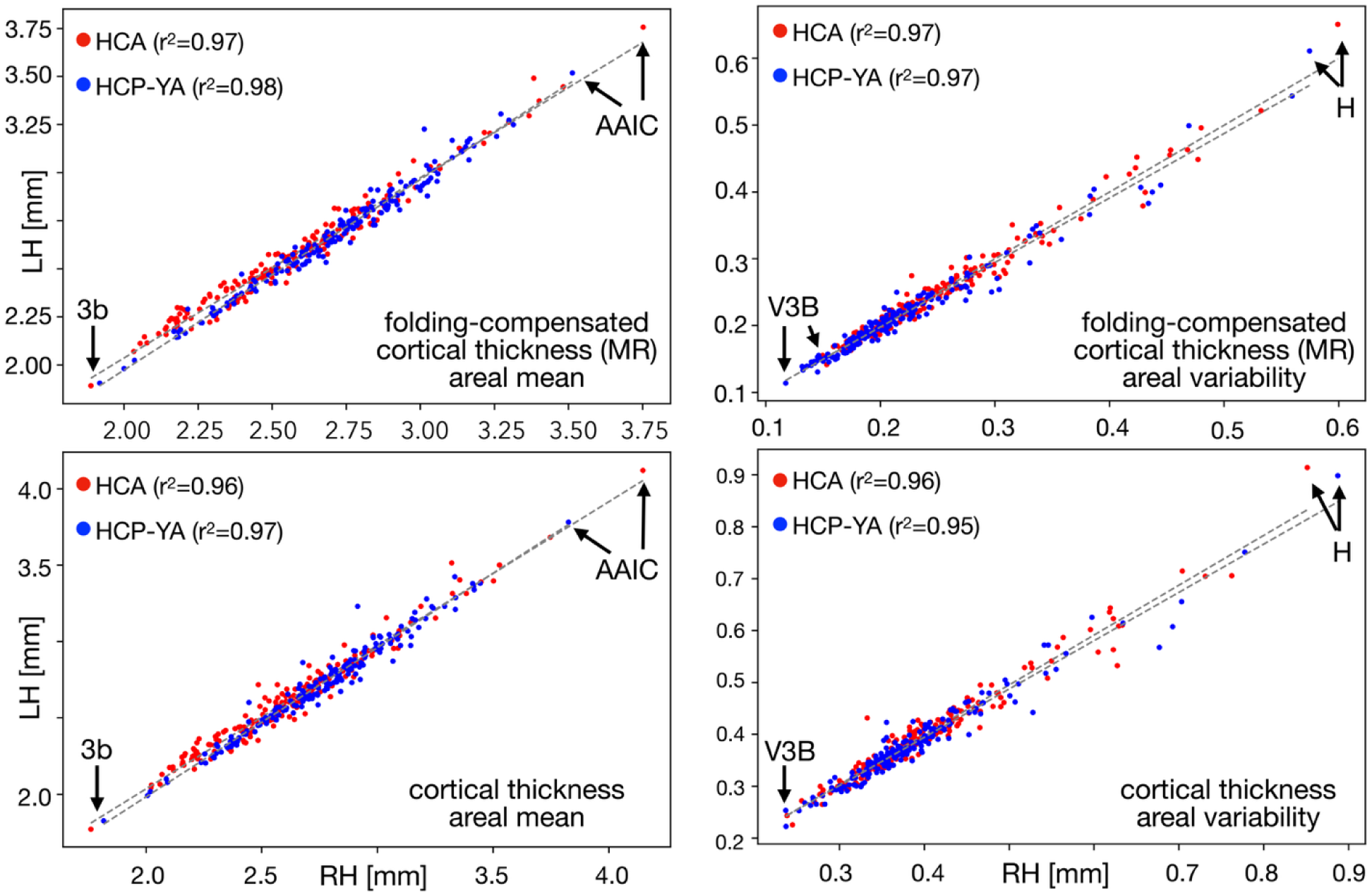
Areal hemispheric symmetry of uncompensated and folding-compensated cortical thicknesses. For both measures the same cortical areas exhibit the minimum (areas 3b and V3B) and maximum values (areas AAIC and H) across the hemispheres (MR: Multiple Regression).

### 3.6 Aging-related changes in cortical thickness and folding-compensated cortical thickness

We examined age-related changes in parcellated mean cortical thickness and folding- compensated cortical thickness estimated using multiple regression across the HCA cohort to determine which measure provides greater explanatory power for aging-related cortical changes (Fig. 13). Our goal was to assess which metric more sensitively and powerfully captures these changes. We performed linear regression analyses between age- and folding-compensated cortical thickness (Fig. 13A) and uncompensated cortical thickness (Fig. 13B) for each area. We then computed the coefficient of determination (𝑅^2^) to quantify the proportion of variance in cortical thickness explained by age-related changes (Fig. 14A), and we used Cohen’s effect size (𝑓^2^) = 𝑅^2^⁄(1 − 𝑅^2^) (i.e., the variance explained by the regression divided by the variance not explained by the regression) to reveal the influence of age on both measures across the cortex. Following conventional interpretations, we considered effect sizes small if 𝑓^2^ < 0.10, and large if 𝑓^2^ ≥ 0.35 (Selya et al., 2012).

**Figure 13:**
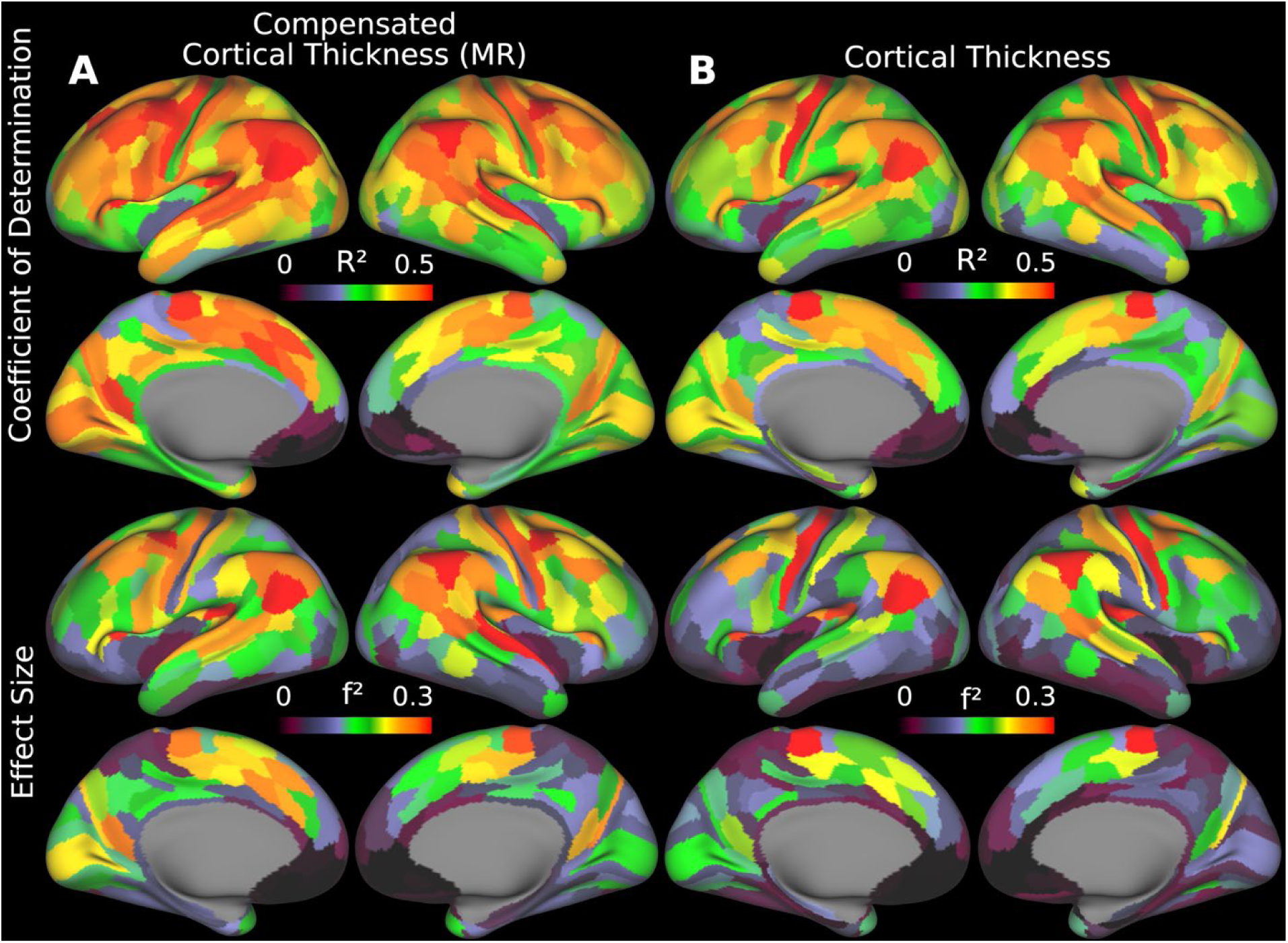
Areal coefficient of determination (𝑅²) and Cohen’s effect size (𝑓²) maps, showing the proportion of variance in (A) compensated and (B) uncompensated cortical thickness explained by age, as estimated using linear regression (MR: Multiple Regression). Data are available at http://balsa.wustl.edu/0rp56.

**Figure 14:**
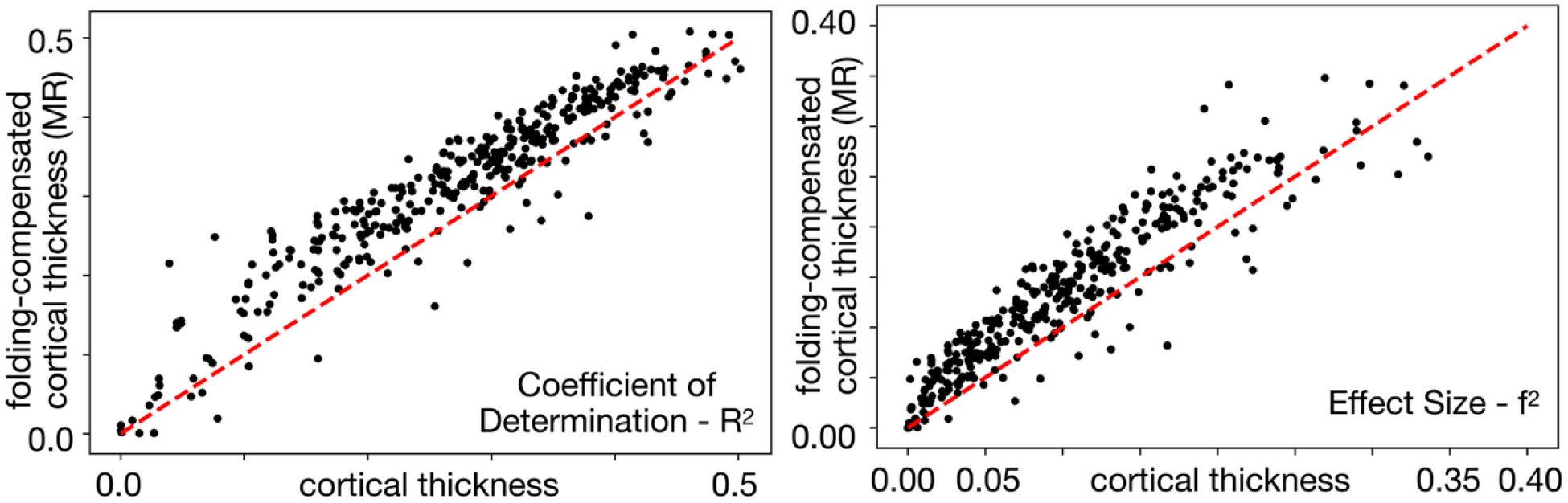
For a large majority of cortical areas, folding-compensated cortical thickness exhibited higher coefficients of determination (𝑅²) and effect sizes (𝑓²), indicating greater explanatory power compared to uncompensated cortical thickness. Red dashed lines show the line of unity (MR: Multiple Regression).

For the majority of areas, the folding-compensated cortical thickness has higher 𝑅*^2^* and 𝑓*^2^* compared to uncompensated cortical thickness (Fig. 14), which indicates that folding- compensated cortical thickness better captures age-related cortical changes by explaining a greater proportion of variance and by demonstrating a stronger effect size of aging across the cortex. The effect size increase is 41% when averaged across all areas weighted by the surface area of each area. Thus, folding-compensated cortical thickness is a more sensitive marker of cortical atrophy across most of the cortex. On the other hand, for some areas that are indicated by black borders in Fig. 15, cortical thickness has higher 𝑅*^2^* and 𝑓*^2^* compared to folding- compensated cortical thickness. Notably, these areas tend to be in locations where cortical folding variability across individuals is less (around the central sulcus, within the insula, and around the medial wall).

**Figure 15:**
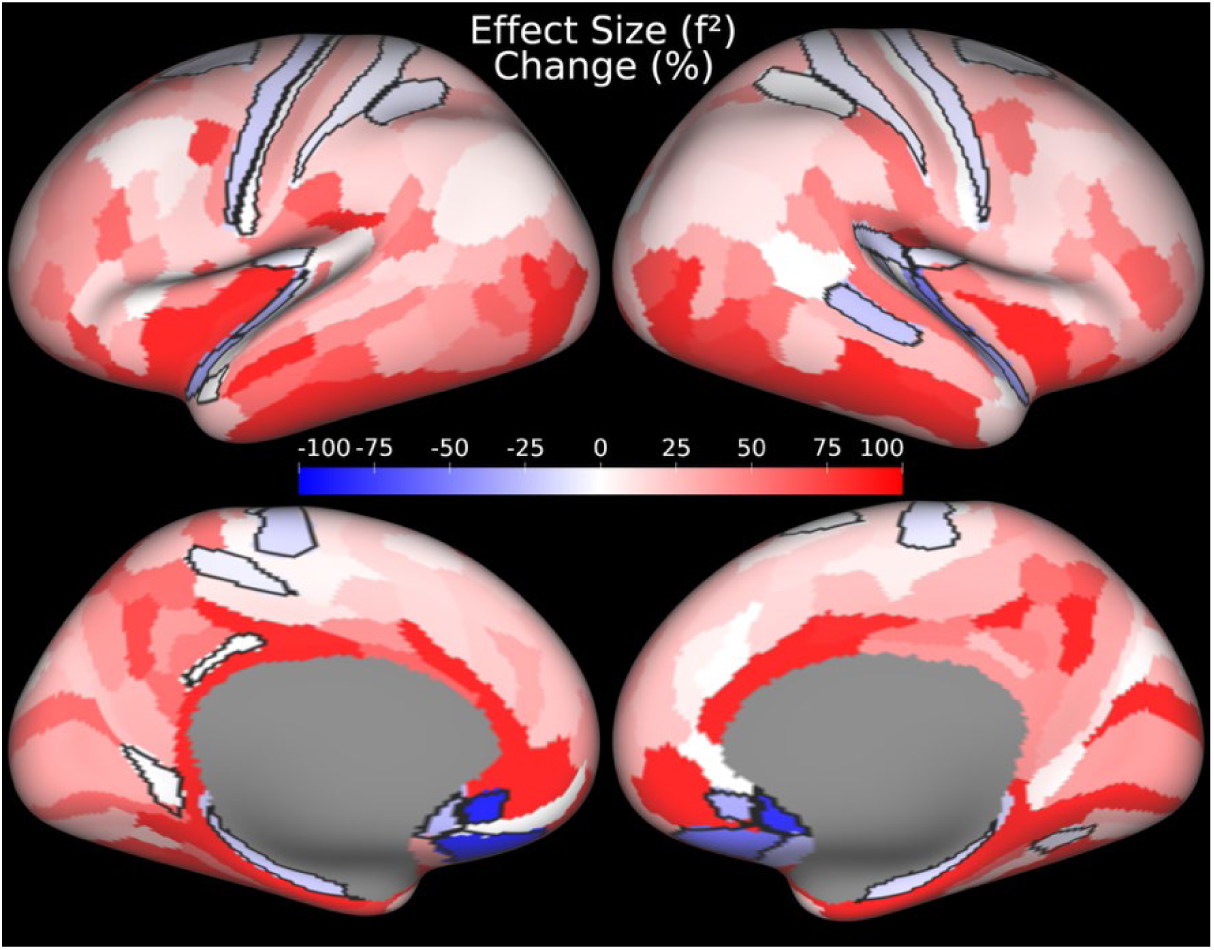
Effect size percentage change between uncompensated cortical thickness and nonlinear folding-compensated cortical thickness with multiple regression. Areas outlined in black indicate areas where the effect size of uncompensated cortical thickness exceeds that of nonlinear folding-compensated cortical thickness. Data are available at http://balsa.wustl.edu/rgMGq.

### 3.7 Spatial gradients defining cortical area borders

We further examined the spatial gradients of local nonlinear folding-compensated cortical thickness and compared them to the gradients of global linear folding-compensated cortical thickness and uncompensated cortical thickness in the group-average HCP-YA cohort. Spatial gradients serve as important markers of cortical area boundaries, where sharp transitions in cortical properties indicate potential areal borders. (Computational details of vertex-wise spatial gradients are provided in detail in Glasser et al., (2016)). Here, we assess whether the newly derived nonlinear folding-compensated cortical thickness exhibits stronger and more distinct gradients that align more closely with other cortical areal features—such as myelin and resting-state functional MRI (rfMRI) gradients—compared to both global linear folding-compensated cortical thickness and uncompensated cortical thickness gradients. A stronger correspondence with these independent modalities would suggest that nonlinear compensation would enhance the delineation of cortical areal borders, potentially aiding our understanding of cortical organization.

Figure 16 illustrates the potential areal borders (white dots) on the group average map of nonlinear folding-compensated cortical thickness (Fig. 16A) and the spatial gradients for nonlinearly compensated (Fig. 16B), linearly compensated (Fig. 16C), and uncompensated cortical thicknesses (Fig. 16D), as well as the gradients of rfMRI (Fig. 16E) and myelin (Fig. 16F). The white and pink arrows highlight two example locations on the cortex where the new measure exhibits strong gradient ridges that closely align with the gradient ridges of rfMRI. In contrast, these ridges are not well captured by either uncompensated cortical thickness or global linear folding-compensated cortical thickness. Similarly, the blue arrow marks another cortical location where local nonlinear folding-compensated cortical thickness demonstrates strong gradient ridges that correspond well with myelin gradients—a pattern that is not observed in either the global linear folding-compensated cortical thickness or the uncompensated cortical thickness.

**Figure 16:**
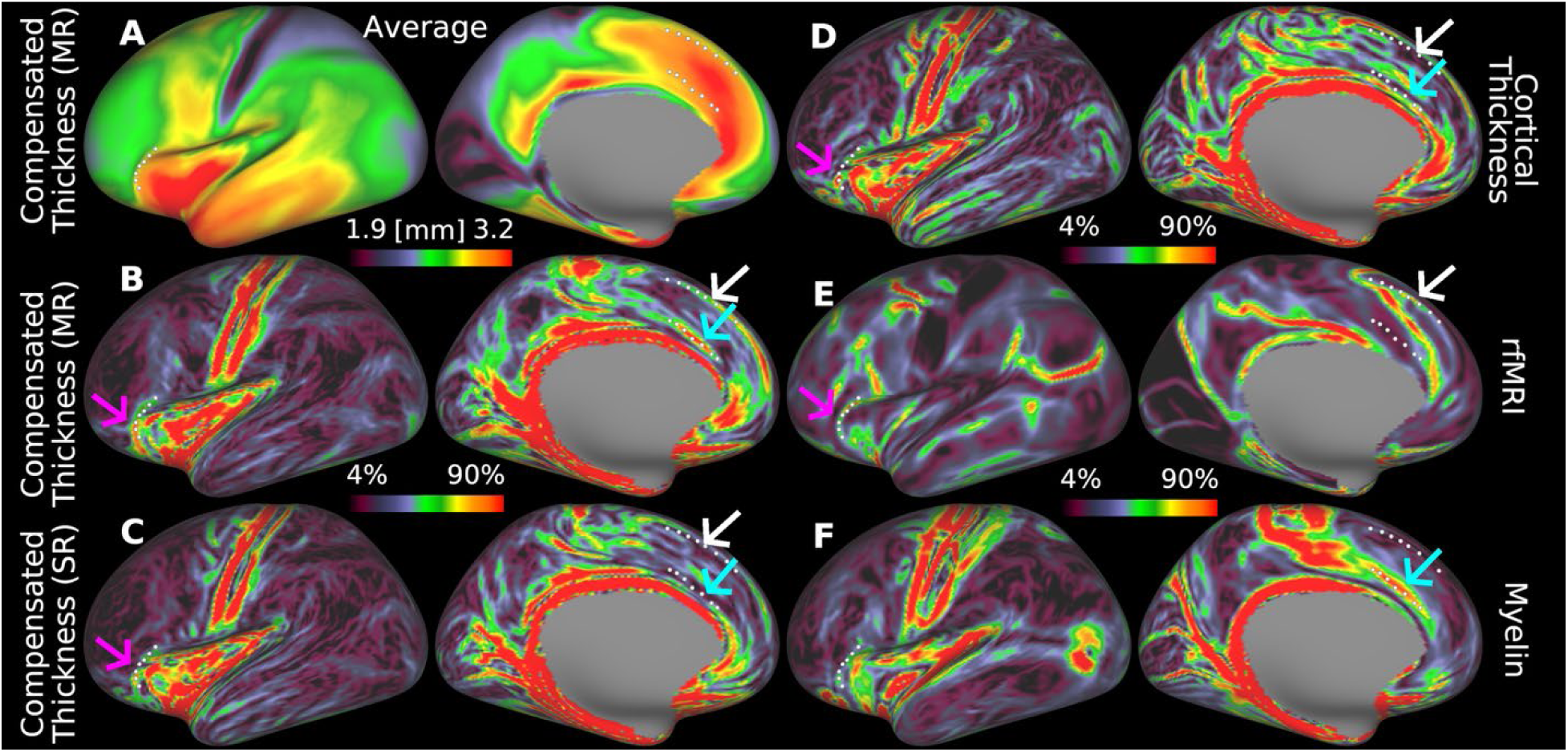
Spatial gradients of multi-modal cortical features. (A) shows the group average local nonlinear folding-compensated thickness, (B) its spatial gradient, and (C-F) spatial gradients of additional cortical features: global linear folding-compensated cortical thickness, uncompensated cortical thickness, resting-state functional MRI (rfMRI), and myelin content, respectively. Local nonlinear folding-compensated thickness shows stronger and more distinct gradient ridges that align closely with gradients in myelin and rfMRI. (SR: Single Regression, MR: Multiple Regression). Data are available at http://balsa.wustl.edu/xv1GN.

Figure 17 illustrates these relationships in the central sulcus, which has a simple and consistent folding pattern: a single long groove with area 4 on the anterior bank, area 3a in the fundus, and area 3b on the posterior bank. Fig. 17A shows a parcellated map of nonlinear folding- compensated cortical thickness with these areas outlined in red. Strings of cyan dots mark key borders: anterior to area 4 (*string a*), 4–3a (*string b*), 3a–3b (*string c*), and posterior to 3b (*string d*). In Fig. 17B, *strings b* and *c* trace either side of the fundus (dark streak) and *strings a* and *d* the gyral crowns (light streaks), confirming that early sensory and motor areas align with folding patterns identified by the mean curvature map. In the uncompensated cortical thickness map (Fig. 17C) and its gradient (Fig. 17D), *string b* coincides with a sharp transition from thick area 4 to thin area 3a, also evident—but more consistently—in the nonlinear folding-compensated thickness and its gradient maps (Fig. 17E, Fig. 17F), noted by the fuchsia arrows pointing to a gap in Fig. 17D that is not present in Fig. 17F. *String b* is modestly expressed in the myelin map (red to orange in Fig. 17G) but strongly in the myelin gradient (red in Fig. 17H). *String b* is accurately delineated along nearly its full length in the nonlinear folding-compensated thickness gradient (Fig. 17D) compared to the uncompensated thickness gradient (Fig. 17D). *String c* is well defined in the myelin gradient (though faint in lateral regions) but is discernible along only a limited strip in Fig. 17F and not at all in Fig. 17D. *String d* is clear in both thickness gradients (Fig. 17D, F), but stronger in the uncompensated thickness map although it is faintly visible in the myelin gradient (Fig. 17H). Finally, *string a* aligns with a strong myelin gradient but there are no corresponding gradient ridges in either thickness maps.

**Figure 17:**
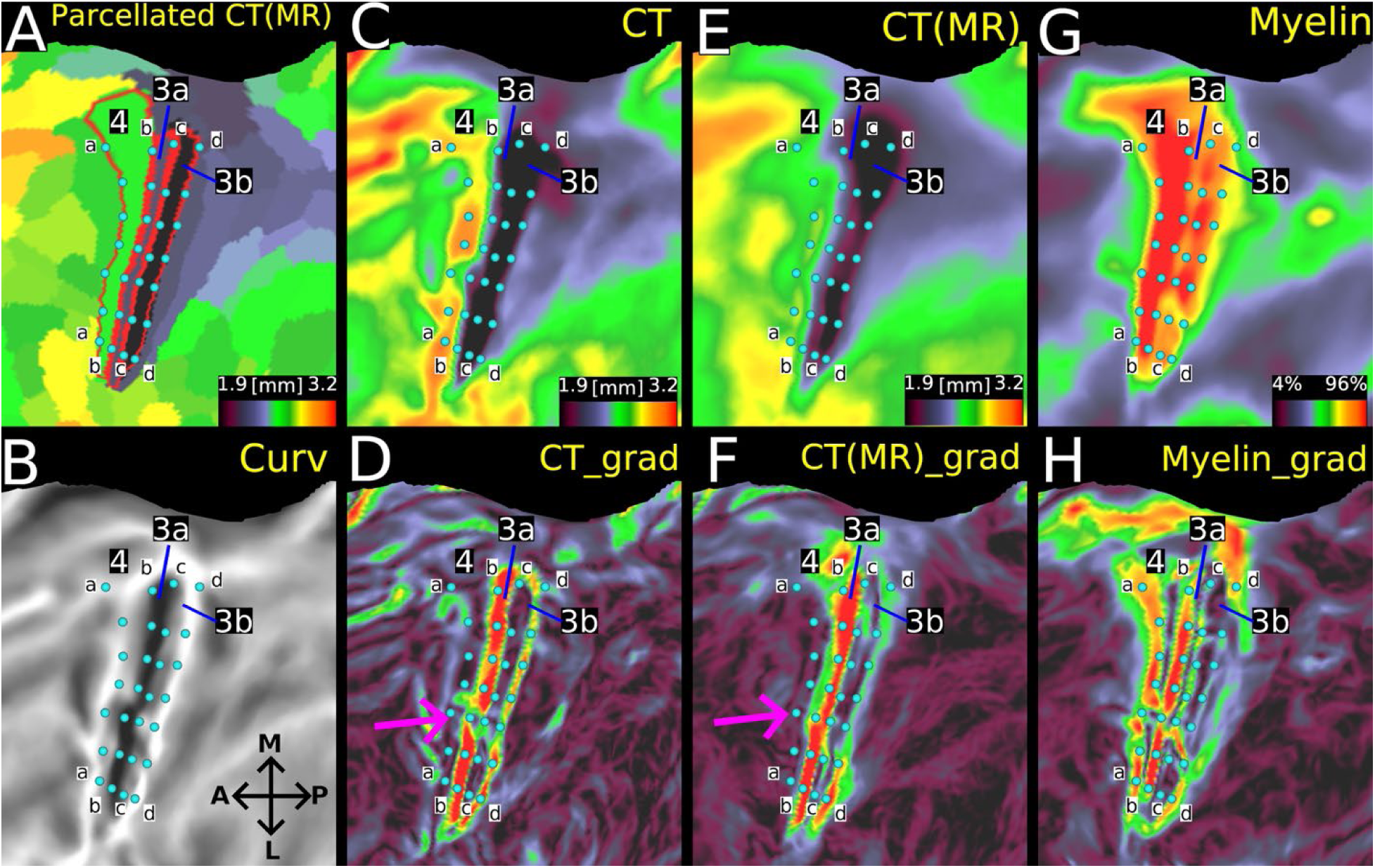
Relationships among nonlinear folding-compensated and uncompensated cortical thickness, myelin, their spatial gradients, and areal borders in the central sulcus of group average HCP-YA data for the MSMAll n=1071 group. (A) shows the parcellated folding-compensated cortical thickness with areal borders for areas 4, 3a, and 3b depicted in red. Areal borders corresponding to transitions between areas are labeled as follows: anterior border of area 4 (*string a*), 4–3a (*string b*), 3a–3b (*string* c), and the posterior boundary of area 3b (*string d*). (B), (C), (E), and (G) show mean curvature, uncompensated cortical thickness, nonlinear folding-compensated cortical thickness, and myelin group-average flat maps of the left hemisphere, respectively. (D), (F), and (H) depict the spatial gradient maps of uncompensated cortical thickness, nonlinear folding-compensated cortical thickness, and myelin respectively. (MR: Multiple Regression, M: Medial, L: Lateral, A: Anterior, P: Posterior). Data are available at http://balsa.wustl.edu/PZwjv.

### 3.8 Folding-compensation versus smoothing of cortical thickness

Many neuroimaging studies apply substantial spatial smoothing to reduce measurement noise, to attempt to account for anatomical variability due to imperfect inter-subject alignment (note that smoothing does not actually improve interindividual alignment, see Coalson et al., (2018)), and to render the data more normally distributed, which typically leads to higher statistical significance values. In published cortical thickness analyses, smoothing is typically performed by linearly filtering thickness maps using surface-based smoothing after resampling to an average cortical surface. However, extensive smoothing markedly reduces the spatial localization fidelity of the results and removes genuine fine-grained spatial patterns. Our review of the literature revealed that Gaussian smoothing of cortical thickness is common (although there are other less commonly used smoothing alternatives, such as heat kernel (Chung et al., 2005) and spherical wavelets (Bernal-Rusiel et al., 2008)), but varies in full-width at half maximum (FWHM) from 10 mm to 60 mm (see Supplementary Table 1). Here, we compared folding-compensation of cortical thickness with Gaussian smoothing and used a FWHM of 24 mm in our analysis, which is the average of commonly used Gaussian smoothing parameters for cortical thickness analyses in the literature. As shown in Figure 17, smoothing at this level considerably reduces the spatial resolution of both the group-average cortical thickness and its gradient, diminishing the visibility of candidate cortical areal boundaries. In contrast, folding-compensated cortical thickness retains much sharper spatial features, preserving fine-scale details in the gradient map.

## 3. Discussion

### 4.1 Overview

We systematically accounted for the effects of cortical folding on cortical thickness in a large cohort of healthy individuals using a local, nonlinear compensation approach based on multiple regression. To robustly correct for folding-induced inter-individual variability, we incorporated five folding measures (maximum and minimum principal curvatures, Gaussian curvature, shape index, and curvedness) into the regression model. Unlike previous approaches, we applied this regression locally, with the optimal patch size determined via a golden-section search optimization algorithm tailored to cortical surface geometry. Our results demonstrate that this locally folding- compensated cortical thickness measure substantially reduces both intra-areal and inter-individual variability, indicating that it is less influenced by folding-related confounds. Importantly, it preserves the same overall pattern of age-related cortical atrophy observed with traditional cortical thickness estimates and thus enhances sensitivity to changes in cortical thickness that may be biologically meaningful. Compared to conventional spatial smoothing, our approach offers superior spatial localization precision. Folding-compensated cortical thickness thus provides a principled alternative to smoothing and holds promise for refining cortical areal parcellation and improving the interpretability of individual differences in cortical thickness. In the following sections, we discuss biomechanical and neurobiological aspects of cortical folding and thickness regulation during cortical development, the efficacy of our compensation method in reducing intra- and inter-individual variability, and its implications for parcellation and future applications in studies of individual differences of cortical thickness.

### 4.2 What regulates cortical thickness and folding during development?

Two mechanisms have been proposed for why the developing cortical sheet remains thin as it expands dramatically in surface area. During early forebrain development, elevated cerebrospinal fluid (CSF) pressure—driven by continuous CSF production from the choroid plexus combined with a diffusion barrier formed by tight junctions between neuroepithelial cells lining the ventricular surface—promotes ventricular expansion and stretches the surrounding cerebral wall, keeping it thin, much like the expansion of a balloon (Van Essen, 2020). In addition, an anisotropy in cortical fiber orientations, particularly the radial alignment of apical dendrites of pyramidal cells, radial glial processes, and developing axons (Wang et al., 2017), may lead to greater mechanical stiffness along the radial axis compared to the tangential plane. This stiffness anisotropy would favor tangential expansion of the cortical sheet (Van Essen, 2023, 2020, 1997). Later in fetal development, increased dendritic arborization in tangential and oblique directions may contribute to the overall thickening of the cortex. Furthermore, regional differences in the degree of stiffness anisotropy could help explain inter-areal variations in cortical thickness (e.g., thick area 4 versus thin area 3b).

Cortical folding, or gyrification—the formation of gyri and sulci—arises from a complex interplay of mechanical and cellular processes and has been extensively studied. From a mechanical perspective, cortical folding is an instability problem of constrained differential growth in a multi-layered system. Buckling can occur as a result of differential expansion of the cortical sheet with respect to the underlying gray/white interface (Akula et al., 2023; Budday et al., 2015a; Kroenke and Bayly, 2018; Richman et al., 1975). Another perspective emphasizes pathway-specific axonal tension that drives folding in specific locations combined with tethering tension that drives buckling or irregular folding (Van Essen, 2023, 2020, 1997). A third perspective posits that differential proliferation and/or migration of cortical progenitor cells in the tangential plane, particularly in the outer subventricular zone that gives rise to superficial cortical layers. The density of progenitor cells varies according to the overall degree of regional cortical expansion. In ferrets, differential proliferation is evident for regions that become a specific gyrus or sulcus (Reillo et al., 2011). In primate cortex, it remains unclear how closely the differential proliferation profile matches the eventual pattern of gyri and sulci (Kriegstein et al., 2006; Lukaszewicz et al., 2006; Van Essen, 2007). Altogether, cortical folding likely involves multiple biomechanical and biological mechanisms acting in concert. Additionally, the relative importance of different mechanisms likely varies across species. In humans, these mechanical and biological mechanisms contribute to a high degree of folding-induced inter-individual variability in cortical morphology (Van Essen et al., 2019). Such variability is especially pronounced in regions with complex tertiary and secondary folds, where subtle differences in local geometry and tissue properties lead to highly individualized cortical patterns (Budday et al., 2015b; Demirci et al., 2023b; Voorhies et al., 2021).

Cortical folding is associated with systematic variations in cortical thickness, with gyri appearing consistently thicker and sulci thinner. Analytical, experimental, and computational models show that the physical forces that emerge during the buckling of a thin elastic sheet on a softer substrate naturally lead to the formation of thick peaks and thin valleys (Holland et al., 2020, 2018). However, mechanical models assuming homogeneous growth across the cortical sheet fall short in replicating the thickness differences between gyri and sulci observed in human MRI data. In contrast, heterogeneous growth models, which incorporate regionally biased expansion— particularly favoring gyral zones—more accurately reproduce the thickness inhomogeneities evident in vivo (Wang et al., 2021). Further, the white matter immediately subjacent to sulcal fundi is dominated by axons running tangential to the gray/white boundary, many of which are so-called ‘U-fibers’ extending to neighboring gyri (Reveley et al., 2015). If these curved axons are under tension, they might tend to compress the overlying sulcal fundus, making it thinner.

Another fundamental question concerns the extent to which the human cerebral cortex conforms to Bok’s equi-volume principle (Bok, 1929; Van Essen and Maunsell, 1980). This principle posits that cortical folding induces minimal changes in local cortical volume, whether in gyral or sulcal regions (hence, *equi-volume*), and preserves the underlying cortical circuitry—such as the number of neurons, dendrites, and synapses—even though cellular shapes may become distorted. Deviations from the equi-volume principle would imply that gyral and sulcal regions differ functionally in their microcircuitry. In ferrets, for example, the ratio of superficial to deep layer neurons varies between gyri and sulci (Reillo et al., 2011; Smart and Mcsherry, 1986). However, studies in humans support the equi-volume principle (Waehnert et al., 2014; Wagstyl et al., 2018). Thus, in the human brain, cortical folding appears to enable compact wiring without introducing large circuitry differences between gyri and sulci.

We propose that folding-compensated thickness provides an estimate of *intrinsic cortical thickness*—that is, how thick the cortex would be in the absence of extrinsic mechanical forces that contribute to cortical folding and buckling. By mitigating these effects, folding-compensated thickness may serve as a more biologically interpretable measure of cortical architecture—less confounded by mechanical distortions imposed by cortical folding.

### 4.3 Folding-compensated cortical thickness reduces cortical folding-related inter-individual variability

Using our novel, locally folding-compensated cortical thickness measure, we demonstrated reduced intra-individual variability across the cortex (Fig. 6) and lower inter-individual coefficient of variation in both the young-adult and aging cohorts (Fig. 8). Compared to uncompensated and globally compensated estimates, the locally compensated maps are substantially more homogeneous and spatially uniform at both the individual and group level (Figs. 6, 7, 8). This homogeneity is evident within cortical areas both visually and quantitatively, with minimized intra-regional variation (Figs. 10, 11) while preserving biologically meaningful cortical thickness patterns, e.g., sharply differentiated thick motor and thin somatosensory cortices (Fig. 17). This finding has neurobiological importance, as altered thickness in these regions may serve as biomarkers of neurodevelopmental or neurodegenerative disorders, such as motor cortex thinning in amyotrophic lateral sclerosis (Mezzapesa et al., 2013), and somatosensory abnormalities in autism spectrum disorders (Ong and Fan, 2023).

We also showed that folding-compensated cortical thickness retains strong bilateral symmetry across cortical areas in both mean and standard deviation (Fig. 12), comparable to that of the uncompensated thickness maps, thereby maintaining the intrinsic anatomical organization of the cortex (Van Essen et al., 2012). Although this bilateral consistency underscores the robustness of our compensation approach, further work is required to investigate whether reported systematic hemispheric asymmetries of cortical thickness reflecting sex differences (Luders et al., 2006), functional specialization with maturity (Zhou et al., 2013), functional connectivity (Liao et al., 2023), and lateralization in language and visuospatial processing (Kong et al., 2018) remain in precisely aligned cortical thickness data measured across thousands of individuals.

### 4.4 Folding-compensated cortical thickness is more sensitive to age-related cortical atrophy

We assessed the cortical thinning related to aging in healthy individuals aged 36–103 years from the HCP-Aging cohort. Our results demonstrate that nonlinear, locally compensated cortical thickness provides a technically superior measure with less folding-induced individual variability (Fig. 8) while retaining highly similar patterns of widespread age-related atrophy as uncompensated cortical thickness (Fig. 9). While cortex-wide thinning with age was evident, the most pronounced atrophy occurred in the medial and dorsolateral prefrontal cortices, association cortex, and dorsal anterior cingulate, consistent with previous findings (Hurtz et al., 2014; Salat, 2004; Shaw et al., 2016). In contrast, minimal thinning was observed in visual, somatosensory, and insular cortices, also in line with prior reports (Shaw et al., 2016). Importantly, the observation that folding-compensated cortical thickness reveals similar patterns of age-related atrophy as traditional cortical thickness is critical, given that cortical thinning is a biomarker for Alzheimer’s and other dementia-related neurodegenerative disorders (Lerch et al., 2005; Pettigrew et al., 2016). We further found that locally compensated cortical thickness showed higher coefficients of determination (𝑅²) and larger effect sizes (𝑓²) in the majority of cortical areas, with an overall 41% increase in effect size, demonstrating improved sensitivity to age-related cortical atrophy (Fig. 14- 15). Thus, this measure is more sensitive to biologically meaningful changes in cortical thickness because nuisance effects of folding variability have been attenuated. These findings suggest that removing folding-related confounds yields a cleaner measure of cortical atrophy and a more powerful metric for studying healthy aging. Areas where traditional cortical thickness outperformed compensated thickness were predominantly regions with low folding-induced individual variability (Fig. 15) (e.g., the central sulcus and regions near the medial wall and in the insula), highlighting the regional specificity of folding-thickness interactions across individuals.

### 4.5 Folding-compensated cortical thickness has the potential to refine cortical areal parcellation and improve human cortical mapping

Cortical area parcellation using spatial gradients is a data-driven approach to delineating functionally and structurally distinct regions of the cortex. Rather than relying solely on anatomical landmarks, this method captures discrete boundaries in brain organization based on features like function, architecture, connectivity, and topography (Felleman and Van Essen, 1991; Glasser et al., 2016; Glasser and Van Essen, 2011; Van Essen and Glasser, 2018).

In this work, we computed spatial gradients as described in Glasser et al., (2016) by following a local plane-fitting approach based on linear least squares regression. Our observations show that spatial gradients derived from folding-compensated cortical thickness align more closely with multi-modal cortical features, including resting-state functional connectivity gradients and myelin maps, in multiple locations (Fig. 16), including in the central sulcus (Fig. 17). In several cortical regions, the folding-compensated thickness gradient reveals stronger gradient ridges, which may correspond to more distinct areal boundaries compared to both uncompensated and globally compensated thickness maps. These findings suggest that folding-compensated cortical thickness may be a valuable marker for refining cortical parcellation and improving multi-modal cortical mapping.

### 4.6 Folding-compensated cortical thickness is an alternative to smoothed cortical thickness in neuroimaging analyses

We also evaluated the impact of spatial smoothing, a common preprocessing step in cortical thickness analyses, on the anatomical precision of cortical thickness measurements. Although smoothing is assumed to reduce inter-individual variability, along with increasing signal-to-noise ratio and data normality (Chung et al., 2005; Han et al., 2006; Lerch and Evans, 2005), it unavoidably blurs fine spatial detail, such as cortical areal boundaries, and compromises the spatial localization of neuroimaging results (Coalson et al., 2018). There is currently no consensus on the optimal smoothing kernel. Some studies advocate for relatively large kernels (e.g., 30 mm FWHM) (Lerch and Evans, 2005), whereas others suggest smaller kernels (e.g., 6 mm) that are sufficient to enhance measurement reliability without excessive spatial degradation (Han et al., 2006). In practice, however, even modest smoothing levels can degrade spatial resolution and reduce the anatomical fidelity of MRI-based estimates (Coalson et al., 2018) (e.g., perhaps up to 4mm FWHM on the surface before more significant degradation of localization to cortical areas occurs). Our review of the literature (see Supplementary Table 1) suggests that smoothing parameters may often be selected arbitrarily, with kernel sizes ranging from 10 mm to as high as 60 mm FWHM, underscoring the lack of standardized practice. Our results demonstrate that applying the mean of the commonly used Gaussian smoothing kernels (24 mm FWHM) to traditional cortical thickness maps blurs spatial gradients and obscures anatomically meaningful areal boundaries (Fig. 18). In contrast, folding-compensated cortical thickness preserves sharp transitions and retains spatial detail. Based on these findings, we recommend the use of folding- compensated cortical thickness in lieu of aggressive spatial smoothing to reduce nuisance folding variability, enhance statistical sensitivity, and maintain the anatomical precision necessary for robust cortical thickness analysis.

**Figure 18:**
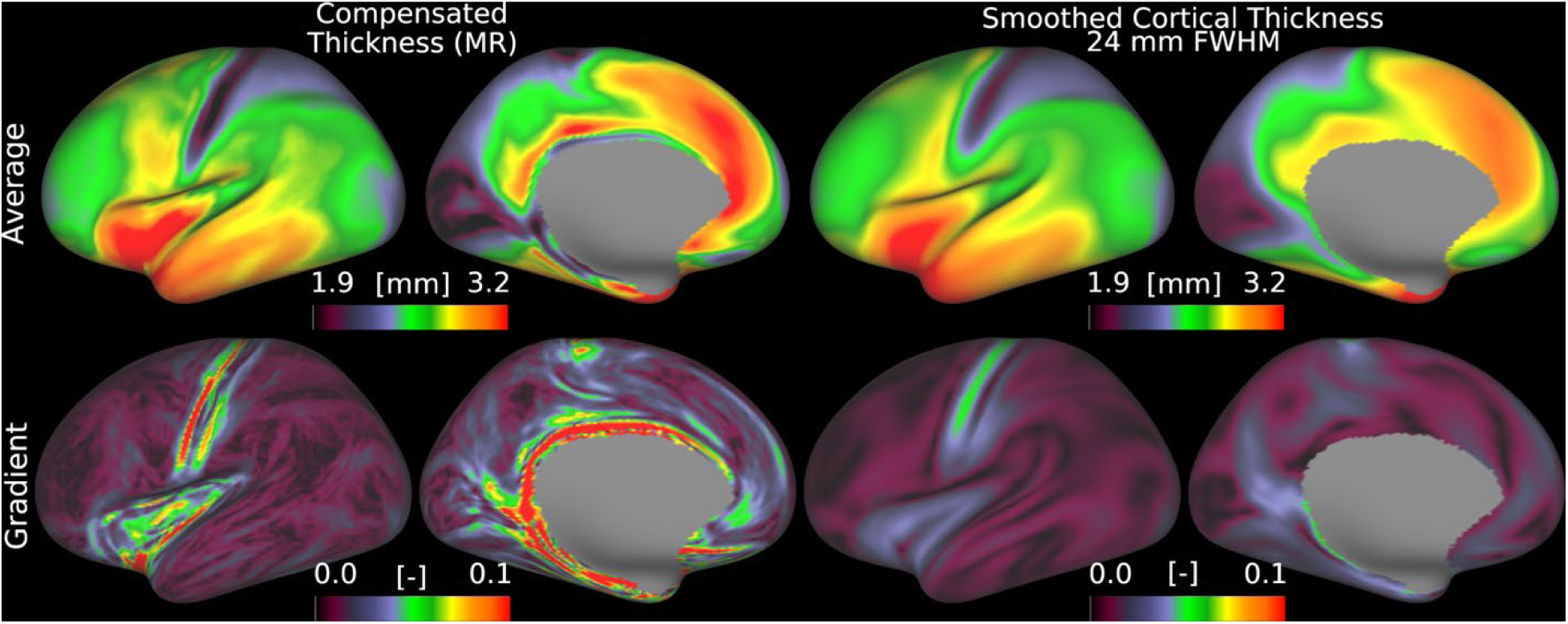
Smoothing versus folding compensation of cortical thickness. Folding-compensation preserves sharp spatial features and fine-scale details in the resulting gradient maps, which are diminished by conventional smoothing (MR: Multiple Regression, FWHM: Full-width at Half Maximum). Data are available at http://balsa.wustl.edu/7X45M.

Collectively, these results highlight the utility of nonlinear, locally adaptive compensation for cortical folding in enhancing the interpretability, sensitivity, and spatial specificity of cortical thickness measurements. By better isolating the intrinsic component of cortical thickness, this approach offers a promising tool for investigating the neurobiological underpinnings of cortical development, individual variability, and age-related neurodegeneration. Furthermore, it may serve as a more sensitive biomarker for neurological conditions in which cortical thickness is known to be altered, including autism spectrum disorder (Khundrakpam et al., 2017), Alzheimer’s Disease (Phan et al., 2024), multiple sclerosis (Fujimori et al., 2021), and age-related atrophy (Hurtz et al., 2014; Salat et al., 2004), as well as psychiatric disorders such as schizophrenia (Kim et al., 2012).

### 4.7 Limitations, alternatives, and future directions

Despite its advantages, our approach has some limitations. First, the accuracy of surface-based thickness estimation is limited by errors in pial and white surface placement, which may contribute to residual high-frequency variability, particularly in regions with thin or highly myelinated cortex (e.g., somatosensory or visual cortex). Improvements in surface placement will likely require both algorithmic improvements (e.g., in FreeSurfer) and increases in spatial resolution of the underlying T1w and T2w (or FLAIR) images (e.g., beyond the 0.7 mm-0.8 mm isotropic MRI scans used here).

Second, although we used a simple cross-sectional design in this study to demonstrate the value of compensated cortical thickness, it is readily applicable to a longitudinal design, which would offer greater insight into the trajectories of gradual, age-associated structural changes in the aging brain. Longitudinal data are particularly valuable for disentangling intra-individual changes from inter-individual variability, and for capturing the nuanced progression of cortical atrophy over time. Notably, prior research has shown that cross-sectional designs may underestimate age-related declines in cortical thickness (Shaw et al., 2016), further highlighting the need for longitudinal studies to more accurately characterize the aging process.

We used mid-thickness surfaces in all our analyses. A potential alternative would be to use mid- equivolume surfaces based on Bok’s equivolume principle, as they more closely approximate laminar architecture and follow the curvature of cortical layers. While our method is applicable in principle to mid-equivolume surfaces, they might be arguably less suitable for cortical thickness compensation. The mid-equivolume surface coordinates depend on the curvature of the white and pial surfaces, which may make them more sensitive to localized white or pial surface placement errors. This may propagate the effects of such placement errors into the subsequent curvature measures more than with the mid-thickness surfaces (Gülban and Huber, 2025).

Furthermore, we selected five curvature measures because they can be defined at local coordinates and directly characterize cortical folding geometry. Alternative measures such as the gyrification index or fractal dimensionality could, in principle, be included, but these are typically computed at the global level, and their local estimates depend on arbitrarily defined surface patches (regions of interest - ROI) around each vertex. Since our compensation method operates on vertices rather than ROIs, we prioritized curvature descriptors that capture folding–thickness dependencies at a vertex-wise scale.

Our analyses focused on healthy young and aging adult cohorts; however, the methodology we introduce could be applied to many clinical populations in future work. For example, our compensation approach may be useful in conditions such as Alzheimer’s disease or autism spectrum disorder, where the relationship between cortical folding and cortical thickness falls within the range observed in healthy human brains (i.e., thin, tightly folded visual cortex to thick, coarsely folded medial prefrontal and anterior temporal cortex). However, for disorders with markedly altered folding (e.g., polymicrogyria or pachygyria-lissencephaly and hemimegaloencephaly), the folding–thickness relationship may differ, and careful revalidation of the model would be important. Similarly, major alterations of cortical folding during development, (e.g., due to in-utero hydrocephalus) would require careful validation.

Finally, folding-compensated cortical thickness could also offer benefits to studies exploring developmental trajectories or comparing cortical architecture across species.

## 4. Conclusion

In this study, we introduced a novel, nonlinear, locally adaptive approach for compensating cortical folding effects on cortical thickness measurements. By incorporating five folding measures into a locally optimized regression framework, our method effectively reduces both intra-areal and inter-individual variability in cortical thickness that is attributable to cortical folding. This folding- compensated cortical thickness measure retains biologically meaningful patterns of cortical organization and age-related atrophy, while significantly enhancing anatomical specificity compared to conventional smoothing techniques.

Our findings demonstrate that folding-compensated thickness offers a principled alternative to spatial smoothing, preserves bilateral anatomical organization, and improves alignment with multi-modal cortical features, including functional connectivity and myelination gradients. These improvements support the utility of this method in refining cortical parcellation, improving individual-level morphological analyses, and enhancing the sensitivity of thickness-based biomarkers in both healthy and clinical populations.

Our results are robust across large cross-sectional datasets of healthy individuals, and future work should use folding-compensated cortical thickness in longitudinal designs and in clinical or developmental populations. Additionally, its application to comparative and translational neuroscience may yield deeper insights into the neurobiological underpinnings of cortical structure and function across species and lifespan stages. Collectively, this work advances our capacity to more accurately and precisely characterize human cortical morphology (i.e., cortical thickness), thereby enhancing the interpretability and translational potential of cortical thickness as a marker of brain health and disease.

## 5. Acknowledgments

This work is supported by the National Institutes of Health under a Ruth L. Kirschstein National Research Service Award (F32MH140418-01).

Data were provided in part by HCP-YA: the Human Connectome Project, WU-Minn Consortium (Principal Investigators: David Van Essen and Kamil Ugurbil; 1U54MH091657) funded by the 16 NIH Institutes and Centers that support the NIH Blueprint for Neuroscience Research, by the National Institute On Aging of the National Institutes of Health under Award Number U01AG052564, and funds provided by the McDonnell Center for Systems Neuroscience at Washington University. Supported in part by Connectome Coordination Facility (CCF) I: R24MH108315, Connectome Coordination Facility II: 1R24MH122820, the McDonnell Center for Systems Neuroscience at Washington University, and NIH R01MH060974 (DCVE, MFG). MFG was also supported by the Adult Aging Brain Connectome Project (AABC) NIH U19AG073585. Computations were performed using the facilities of the Washington University Research Computing and Informatics Facility, which were partially funded by NIH grants S10OD025200, 1S10RR022984-01A1 and 1S10OD018091-01. MAH acknowledges support from the National Science Foundation (NSF) under Grant no CMMI 2144412.

## 6. Author Contributions

ND, TSC and MFG conceptualized this study and designed the methodology. ND conducted the data analysis, developed the software, and wrote the first draft. TSC, MAH, DVE and MFG contributed to the revisions and editing of the manuscript. DVE and MFG funded and supervised the work. TSC, DVE and MFG provided guidance on data analysis and results interpretation.

## 7. Ethics

This article complies with the ethical guidelines of the MIT Press: Journal Publication Ethics. The research presented has been conducted and reported with integrity, transparency, and accountability. No specific ethical issues or conflicts of interest are present in this submission.

## Supplementary Materials

**Supplementary Figure 1:**
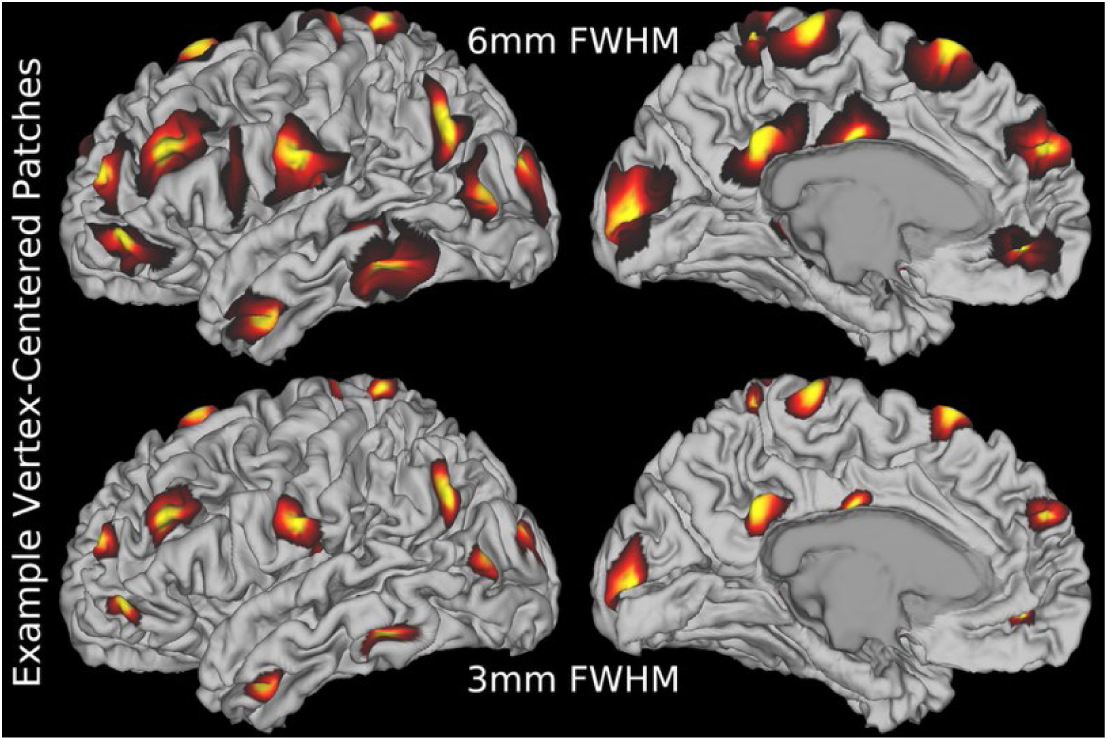
Two example vertex-centered patches within the selected search range for optimal patch size (3–10 mm). The minimum size of 3 mm is sufficient to include at least one inward and one outward fold, which is necessary for capturing local folding geometry relevant to our analysis. Medial wall vertices are masked and excluded from all our analyses. Data are available at http://balsa.wustl.edu/66vrZ.

**Supplementary Figure 2:**
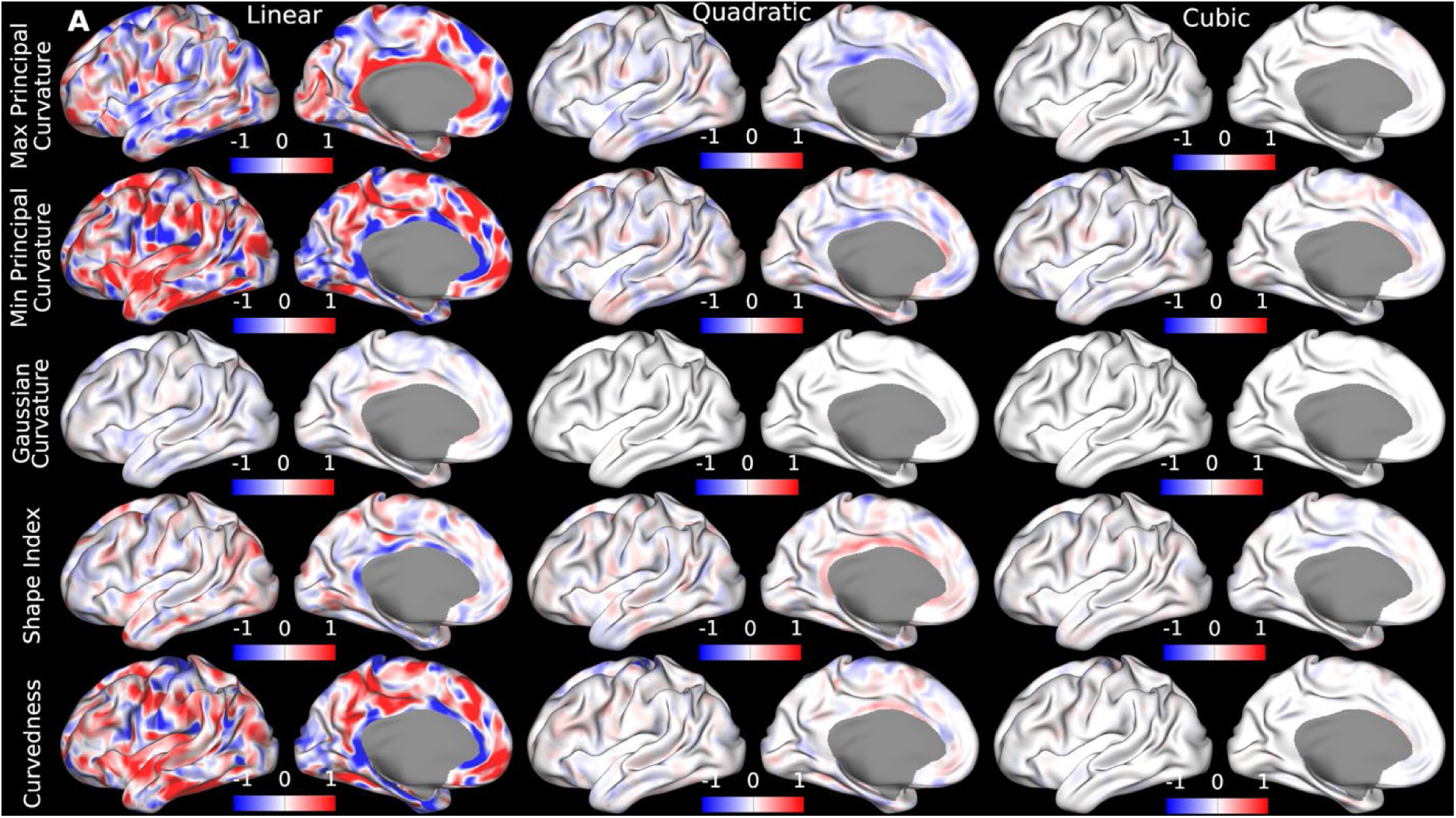
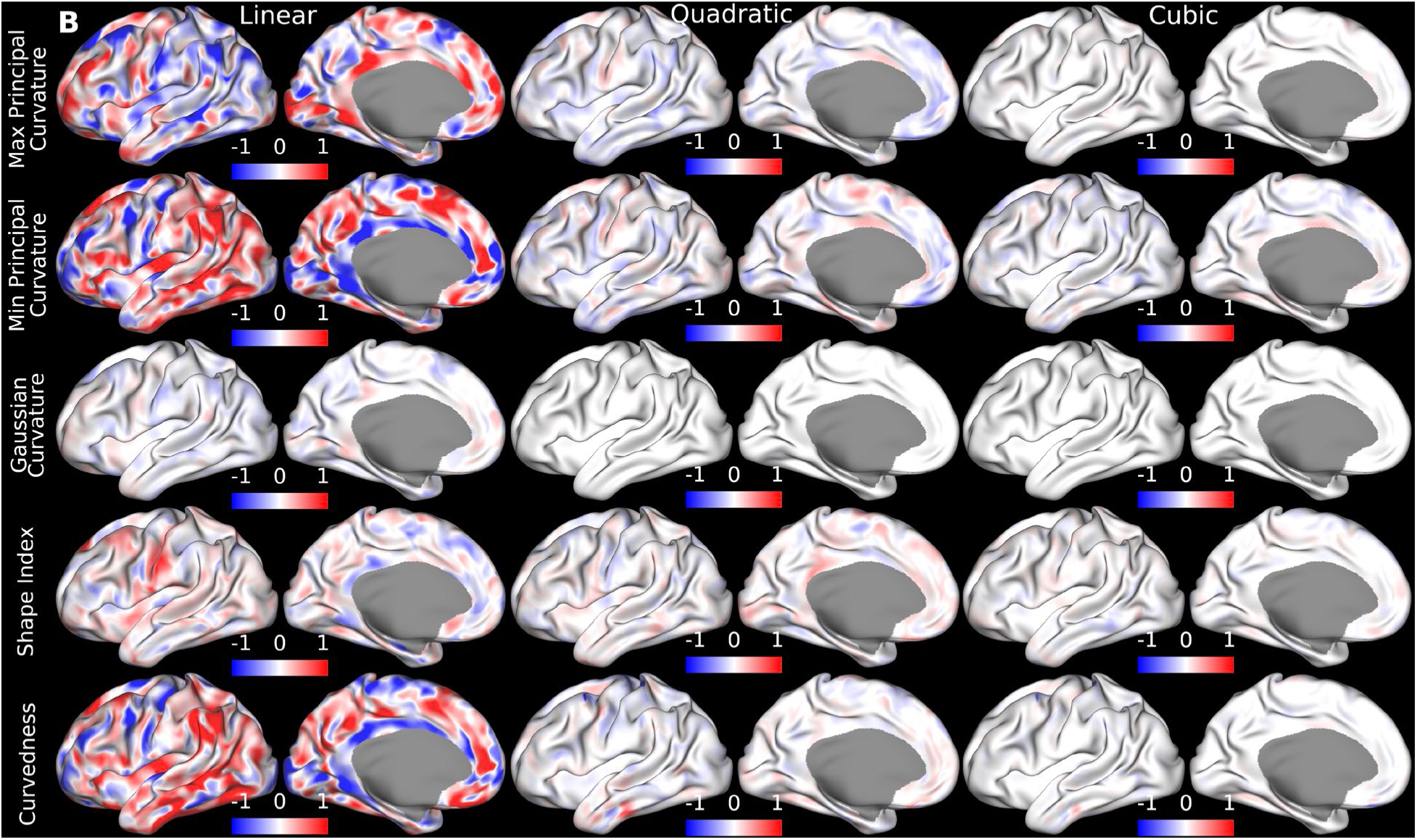
Vertex-wise (normalized with the standard deviation of the corresponding curvature feature) regression coefficient maps displayed on the inflated surface of two individuals of the HCP-YA dataset (A: case 100408, B: case 100307). Red and blue represent positive and negative modeled effect of a curvature feature on thickness, respectively. Cortical thickness is more strongly associated with principal curvatures and curvedness and less strongly with Gaussian curvature, as Gaussian curvature fails to disambiguate between a cup and a cap morphology. Linear regression coefficients show the strongest effects, while quadratic terms contribute moderately and cubic terms minimally. Accordingly, we included terms up to the second order in our polynomial model, as cubic coefficients account for only negligible effects. Data are available at http://balsa.wustl.edu/1r95D and http://balsa.wustl.edu/56rXD.

**Supplementary Table 1:** Summary of full width at half-maximum (FWHM) values used for cortical thickness smoothing in prior studies. This table lists representative smoothing kernel sizes reported in the literature, illustrating the range of values commonly applied in surface-based cortical thickness analyses. In our review, if the smoothing parameter is presented in 𝜎 (standard deviation) we converted it into FWHM for consistency.

| Resource | FWHM (mm) |
| --- | --- |
| (Chung et al., 2005) | 30 |
| (Doyle-Thomas et al., 2013) | 20 |
| (Hurtz et al., 2014) | 10 |
| (Hutton et al., 2008) | 12 |
| (Kim et al., 2012) | 10 |
| (Kuperberg et al., 2003) | 60 |
| (Lerch et al., 2005) | 20 |
| (Misaki et al., 2012) | 15 |
| (Nam et al., 2015) | 20 |
| (Rosas et al., 2002) | 16 |
| (Salat et al., 2004) | 52 |
| (Lerch and Evans, 2005) | 30 |

